# LAG3 regulates antibody responses in a murine model of kidney transplantation

**DOI:** 10.1101/2022.01.31.478518

**Authors:** Michael Nicosia, Ran Fan, Juyeun Lee, Victoria Gorbacheva, José I. Valenzuela, Yosuke Yamamoto, Ashley Beavers, Nina Dvorina, Eduardo Chuluyan, Motoo Araki, William M. Baldwin, Robert L. Fairchild, Booki Min, Anna Valujskikh

## Abstract

Lymphocyte activation gene-3 (LAG3) is a coinhibitory receptor expressed by a range of immune cells. While immunomodulatory potential of LAG3 is being actively explored in cancer and autoimmunity fields, there is no information on how this pathway affects alloreactive immune responses following organ transplantation. The goal of this study was to investigate the functions of recipient LAG3 in a mouse model of renal allograft rejection. We found that mice deficient in LAG3 expression have elevated heterologous immunity against a panel of alloantigens prior to transplantation. Recipient LAG3 deficiency results in rapid rejection of MHC-mismatched renal allografts that are spontaneously accepted by WT recipients, with graft histology characteristic of antibody mediated rejection (ABMR). Depletion of recipient B cells but not CD8^+^ T cells significantly extended kidney allograft survival in LAG3^-/-^ recipients further supporting ABMR as the main mechanism of graft loss. Treatment of WT recipients with an antagonistic LAG3 antibody enhanced anti-donor immune responses and induced kidney damage associated with chronic rejection. The experiments using conditional LAG3 knockout recipients demonstrated that LAG3 expression on either T or B cells is sufficient to regulate anti-donor humoral immunity. These results are the first to identify LAG3 as a regulator of both T and B cell responses to kidney allografts and a potential therapeutic target for ABMR prevention and treatment.

## Introduction

Lymphocyte activation gene 3 (LAG3, CD223) is an immune checkpoint inhibitor of the immunoglobulin superfamily, closely related to CD4^1^. LAG3 is expressed on a number of immune cells including CD4^+^ and CD8^+^ T cells, T regulatory cells, B lymphocytes^1–6^, plasmacytoid dendritic cells^7^ (pDCs) and NK cells^1^. Similar to other coinhibitory receptors such as CTLA4 or PD-1, LAG3 is not expressed by naïve T cells, but its expression is induced on CD4^+^ and CD8^+^ T cells following TCR engagement^1^. Murine LAG3 shares 69.9% sequence homology with the human protein preserving structure and function, and largely mirrors the expression profile of human LAG3^8, 9^. Analogous to CD4, LAG3 binds to major histocompatibility complex II molecules (MHC-II) on antigen presenting cells (APCs)^10, 11^, and additionally interacts with other ligands such as Galectin-3^12^, liver sinusoidal endothelial cell lectin (LSECtin)^13^, α-synuclein fibrils (α-syn)^14^, and fibrinogen-like protein 1 (FGL1)^15^.

In T cells, LAG3 is co-localized with T cell receptor/CD3 complex within immune synapse and inhibits TCR signaling leading to impaired cell activation, proliferation, differentiation, and effector functions in the context of autoimmunity, infection and tumor^16–22^, although the underlying mechanisms remain unclear. It was recently reported that, acidification via a conserved tandem glutamic acid-proline repeat in the LAG3 cytoplasmic tail lowers the local pH at the immune synapse, and dissociates Lck from the CD4 or CD8 co-receptor, resulting in a loss of co-receptor-TCR signaling and limited T cell activation^23^. In addition, LAG3 plays an essential role in T regulatory cells (Treg)^6, 24, 25^, and mice with Treg-specific LAG3 deficiency are resistant to autoimmune diabetes^26^. In contrast, there is a paucity of studies investigating the role of LAG3 in B lymphocytes. Activation of murine splenocytes *in vitro* induces the endogenous expression of LAG3 on CD19^+^ B cells in T cell and soluble factor dependent manner ^3^. However, the functional consequences of LAG3 up-regulation in B cells during humoral immune responses remain to be investigated. LAG3 expression has been shown to be a defining feature of an IL-10 producing subset of plasma cells^27^. While it was demonstrated that these cells arise from BCR dependent signals and that their IL-10 production is dependent upon innate signals such as TLRs, the role of LAG3 expression on this B cell subset is still poorly understood.

Even though the relative importance of LAG3 on conventional T cells, Tregs, B cells and other cell types during immune responses still remains to be determined in different settings it is a molecule of great interest for clinical interventions. A number of monoclonal antibodies targeting LAG3 with either antagonistic, depleting, or agonistic activities have been developed and are currently in clinical trials for cancer and autoimmunity patients^28–37^. However, LAG3 is not commonly considered as a therapeutic target in organ transplantation, largely due to the paucity of information on the role of this pathway in experimental animal models. Lucas *et al*.^38^ reported that LAG3 blockade prevents tolerance induction in alloreactive CD8^+^ T cells in a mouse model of allogeneic bone marrow transplantation, but that this effect did not depend on LAG3 expression by CD8^+^ T lymphocytes themselves. In a different study, the depletion of LAG3^+^ T cells with monoclonal antibody extended rat cardiac allograft survival, yet prevented tolerance induction via donor-specific cell transfusion^39^. These findings suggest LAG3 involvement in different aspects of allograft rejection and tolerance. Whereas the functions of CTLA4 and PD1 coinhibitory molecules in transplantation have been extensively studied, it is unclear whether and how LAG3 regulates alloimmune responses to solid organ transplants.

In the current study, we investigated the contribution of LAG3 to alloimmune responses using a mouse model of life-supporting renal transplantation and found that non-transplanted mice deficient in LAG3 expression have elevated cellular and humoral heterologous immunity against a panel of alloantigens. Recipient LAG3 deficiency resulted in rapid rejection of fully MHC-mismatched renal allografts that are spontaneously accepted by wild type recipients. While LAG3^-/-^ recipients developed increased cellular and humoral anti-donor immune responses, the renal graft tissue injury was characteristic of antibody-mediated rejection and was significantly diminished by the depletion of B cells but not of CD8^+^ T lymphocytes. Neither T cell– or B-cell specific LAG3 conditional knockout recipients rejected renal allografts indicating that LAG3 deficiency in both cell types is required for dysregulated alloantibody production in the context of transplantation. These data present the first evidence for LAG3 involvement in regulation of humoral alloimmune responses and identify LAG3 as a potential target for diminishing pathogenic alloimmunity.

## Materials and Methods

### Animals

Male C3H/HeJ (C3H, H-2^k^), C57BL/6J (B6, H-2^b^), BALB/cJ (Balb/c, H-2^d^) DBA1/J (DBA H-2^q^) and SJL/J-*Pde6b^rd^*^1^ (SJL, H-2^s^), B6.129P2(C)-Cd19 tm1(cre)Cgn/J (CD19Cre), B6.LAG3^-/-^ aged 6-8 week, were purchased from The Jackson Laboratory (Bar Harbor,ME). LAG3^fl/fl^ mice were commissioned by Biocytogen (Beijing, China) and bred to CD4Cre or CD19Cre mice to generate T cell or B cell conditional LAG3^-/-^ mice respectively. All the mouse protocols were approved by the Institutional Animal Care and Use Committee (IACUC) of the Cleveland Clinic. The mice were maintained under specific pathogen-free conditions at the Cleveland Clinic Lerner Research Institute Biological Resource Institute.

### Kidney Transplantation and Graft Evaluation

Transplantation of murine kidneys was performed as previously described^40^. In brief, the kidney and its vascular supply and ureter were harvested *en bloc*, the donor vasculature were anastomosed to the recipient abdominal aorta and inferior vena cava. Excised kidneys were perfused with University of Wisconsin (UW) solution (320mOsm; Preservation Solutions, Elkorn, WI) and stored on ice for 0.5 hours before transplantation. Urinary reconstruction was performed as previously published^41^. The remaining native kidney was removed at the time of transplant, ensuring that survival of the mouse was dependent upon functionality of the transplanted kidney. Graft survival was assessed by daily examination of overall animal health. Recipients were euthanized when low and slow mobility in addition to a hunched posture indicated rejection. At time of euthanasia serum creatinine levels were measured using the VetScan i-STAT1 Analyzer (Abaxis, Union City, CA). Grafts were collected and fixed in methanol before embedding in paraffin for staining with hematoxylin and eosin or Gomori’s trichrome C, and the following antibodies: rabbit monoclonal anti-mouse/human Mac-2 (clone M3/M8; Biolegend #125401), rabbit monoclonal anti-mouse CD4 (clone EPR19514; Abcam #ab183685), rabbit monoclonal anti-mouse CD8 (clone EPR21769; Abcam #ab217344), polyclonal rabbit anti-mouse C4d^42^, rabbit polyclonal anti-Human/mouse vWf (Dako #A0082), rat monoclonal anti-mouse/rat FOXP3 (clone FJK-16s; eBioscience #14-5773) and rat monoclonal anti-mouse B220/CD45R (clone RA3-682; BD Pharmingen #550286). Each stain was evaluated based on two to three complete cross sections from 4-5 grafts per group. Blood urea nitrogen (BUN) levels were measured using BUN Colorimetric Detection Kit (Thermo Fisher Scientific, Waltham, MA). Urine levels of TIM-1/KIM-1/HAVCR and Lipocalin-2/NGAL were measured with Quantikine ELISA kits (R@D System Inc., Minneapolis, MN) as previously published (37084848).

### Recipient Treatment

To deplete CD8^+^ T cells recipients were treated with monoclonal anti-mouse CD8 antibodies, clones TIB105 and YTS169 (Bio X Cell, 0.2mg of each i.p. on days –3, –2, –1 prior to transplantation and every 5 days posttransplant for the duration of the experiment). For B cell depletion, recipients were treated with anti-mouse CD19 clone 1D3 and anti-mouse B220 clone RA3.3A1/6.1 (TIB-146) monoclonal antibodies Bio X Cell, 0.2mg of each i.p. on day 3 and every 5 days posttransplant for the duration of the experiment). For LAG3 blockade, anti-LAG-3 mAb (clone C9B7w, Bio X Cell)^9, 38^ was given i.p. at a dose of 100 μg on days 3, 5, 7, 9, 11, 13 and 15 posttransplant.

### Measurement of Serum Alloantibody

Alloreactive serum antibody titers were measures as previously published^43^. In brief, serum samples were collected via tail-vein bleeding from naïve 10 weeks old mice or from kidney allograft recipients at d. 14 posttransplant and stored at –20°C. Flat-bottomed 96-well plates (Thermo Fisher Scientific) were coated overnight at 4°C with biotinylated class 1 D^d^ (folded with RPGGRAFVTI peptide), class 1 D^k^ (folded with RRLGRTLLL peptide), class 2 I-A^d^ (folded with PVSKMRMATPLLMQA class 2–associated invariant chain peptide), or class 2 I-A^k^ (folded with PVSKMRMATPLLMQA class 2–associated invariant chain peptide) MHC monomers at 1μg/ml in PBS. Peptide/MHC monomers were provided by the National Institutes of Health Tetramer Core Facility at Emory University (Atlanta, GA). Plates were washed with PBS and blockade with 1% BSA-PBS solution for 1h at room temperature. A total of 100μl of serum diluted from 1:100-1:500 were added to the plates and incubated overnight at 4°C. Plates were washed with PBS and PBS-Tween (0.25%), and then incubated with goat anti-mouse IgG –horse radish peroxidase (HRP) conjugate (1:20000 dilution; Thermo Fisher Scientific) or IgG1, IgG2a, IgG2b, IgG3-horse radish peroxidase (HRP) conjugate (1:5000 dilution; Thermo Fisher Scientific) for 1 hour at room temperature. After incubation with conjugate-bound antibodies, plates were washed and developed with ABTS Peroxidase Substrate (KPL, Gaithersburgh, MD). Absorbance was measured at 415nm using the Bio-Rad iMark Microplate Reader (Bio-Rad Laboratories, Hercules, CA).

### ELISPOT Assay

Interferon gamma (IFNγ) ELISPOT assays were performed as previously described using capture and detecting antibody from BD Pharmingen^44^. Spleens from recipients or naïve non-transplanted mice were stimulated with mitomycin C-treated donor C3H or BALB/c, SJL or DBA spleen cells for 24 hours. The resulting spots were analyzed using an ImmunoSpot Series 4 Analyzer (Cellular Technology, Cleveland, OH).

### Flow Cytometry

Fluorochrome conjugated antibodies were purchased from BD Biosciences (San Jose, CA) or eBioscience (San Diego, CA). Cells were isolated from the spleen and stained as previously described^45^, ≥100,000 events per sample were acquired on a BD Biosciences LSRII or a BD Biosciences LSRFortessa X-20 followed by data analysis using FlowJo software (TreeStar, Inc., Ashland, OR).

### Statistical Analyses

Kidney allograft survival was compared between groups by Kaplan-Meier analysis. All other results were analyzed by using a parametric unpaired t-test. The difference between groups was considered significant if the p value was <0.05. Unless noted otherwise, the data are presented as meanL±LSD values. Total numbers of animals in each experimental group are indicated in respective figure legends.

## Results

### Mice deficient in LAG3 have elevated heterologous alloimmune responses prior to transplantation

As LAG3 was demonstrated to play critical role in immune tolerance, we initially evaluated the immune cells from naïve non-transplanted 10 week old B6.WT and B6.LAG3^-/-^ mice. LAG3 deficiency resulted in modest increase in splenic cellularity (data not shown) and in the increased numbers of total and effector memory (TEff_M_) CD4^+^ and CD8^+^ T cells, as well as Tregs compared to WT mice. In addition, mice deficient in LAG3 had elevated numbers of T follicular helper (Tfh) and T follicular regulatory (Tfr) cells in the spleen (**Fig. 1A**, **Fig. S1**). The numbers of major spleen B cell subsets – follicular (FoB), marginal zone (MzB), and transitional (TrB) B cells – were not significantly different between WT and LAG3^-/-^ mice. Notably, mice deficient in LAG3 had significantly increased numbers of germinal center B cells (GcB) and plasma cells (PC) in the spleen suggesting ongoing B cell activation (**Fig. 1B**, **Fig. S2**). There was a trend towards the increase in numbers of regulatory B cells (Bregs) defined as CD19^+^CD1d^hi^CD5^+^ cells. It should be noted that despite the observed shifts towards activated immune cell phenotypes, unmanipulated LAG3^-/-^ mice did not exhibit signs of autoimmune disease up to 6 months of age.

**Figure 1.**
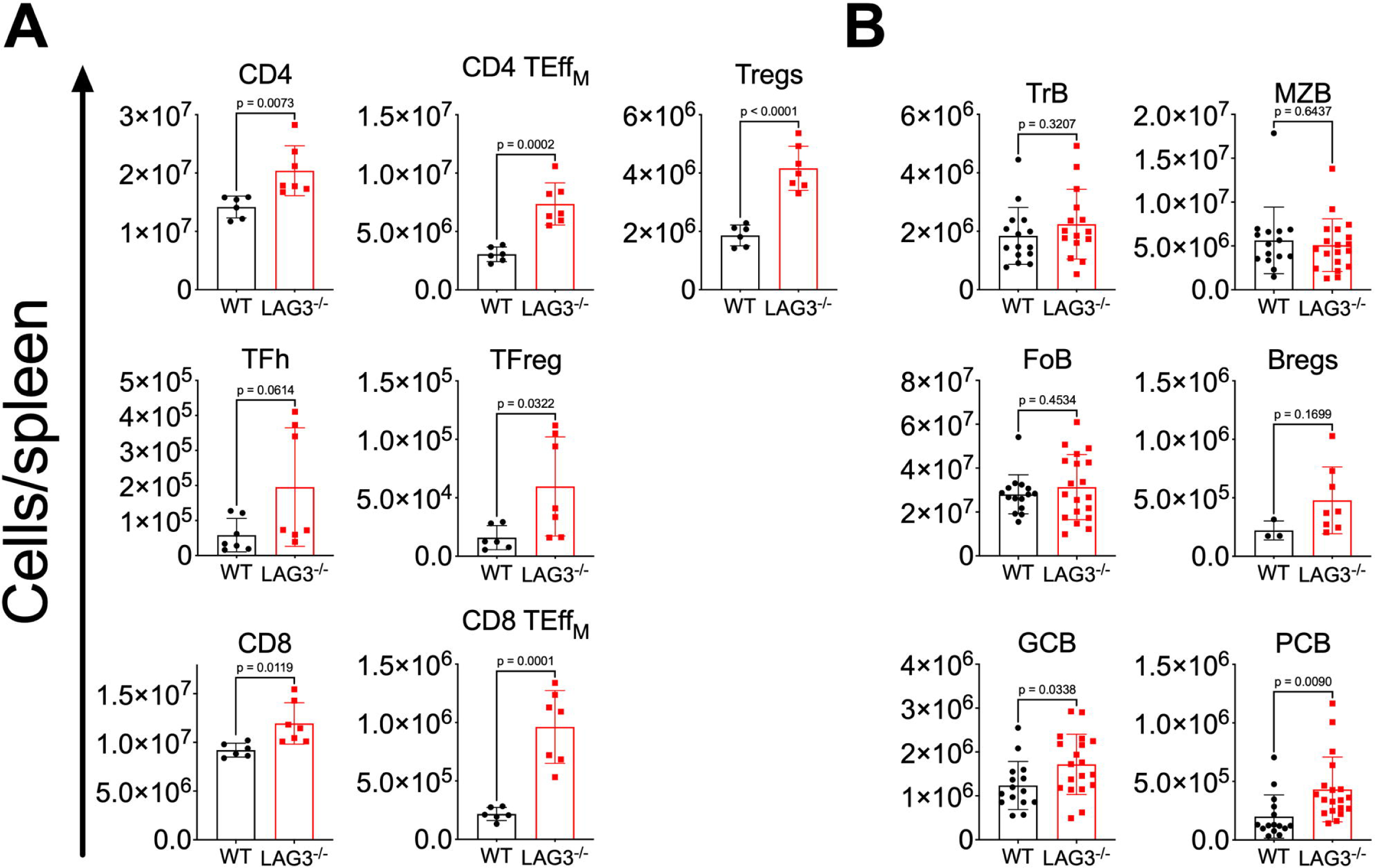
LAG3 deficient mice have expanded lymphocyte subsets. Naïve none-transplanted B6.WT and B6.LAG3^-/-^ mice were sacrificed at 10 weeks of age. **A.** Splenic T cell composition: CD4 (CD3^+^CD4^+^), CD8 (CD3^+^CD8^+^), Tregs (CD3^+^, CD4^+^, FoxP3^+^), CD4 TEff_M_ (CD3^+^, CD4^+^, CD62L^lo^, CD44^hi^), CD8 TEff_M_ (CD3^+^, CD8^+^, CD62L^lo^, CD44^hi^), Tfh (TCRb^+^, CD4^+^, FoxP3^-^, PD-1^+^, CXCR5^+^) and Tfr cells (TCRb^+^, CD4^+^, FoxP3^+^, PD-1^+^, CXCR5^+^). **B.** The composition of splenic B cell subsets. Subsets were defined as follows; FoB –B220^+^, IgM^int^, CD21/35^int^, MZB – B220^+^, IgM^hi^, CD21/35^hi^,TrB – B220^+^, IgM^hi^, CD21/35^lo^, Bregs – CD19^+^, CD1d^hi^, CD5^+^, GCB – B220^+^, GL7^+^, CD38^lo^, and PC –B220^-^, CD138^hi^. The data represents at least two pooled experiments where each symbol represents an individual mouse. Student’s T tests were performed and p<0.05 were considered significant.

To assess the impact of this phenotypic shift on alloimmunity, we probed both B6.WT and B6.LAG3^-/-^ for T cell reactivity against a panel of allogeneic strains (**Fig. 2A**). IFNγ ELISPOT assay demonstrated that compared to WT mice, naïve LAG3^-/-^ mice have increased frequencies of preexisting memory T cells reactive against BALB/c (H2-D^d^), C3H (H2-D^k^), and to a lesser extent, SJL (H-2D^s^) and DBA (H2-D^q^) alloantigens (**Fig. 2A**). We also tested for the presence of alloreactive antibodies in the sera of non-transplanted mice (**Fig. 2B**). Some of the tested naïve LAG3^-/-^ (H2-D^b^) contained elevated levels of IgG antibodies against class I and class II MHC molecules from BALB/c (H2-D^d^, I-A^d^) and C3H mice (H2-D^k^, I-A^k^). These results suggest that LAG3 is an important regulator of immune cell homeostasis and as such may play a crucial role during immune responses to transplanted organs.

**Figure 2.**
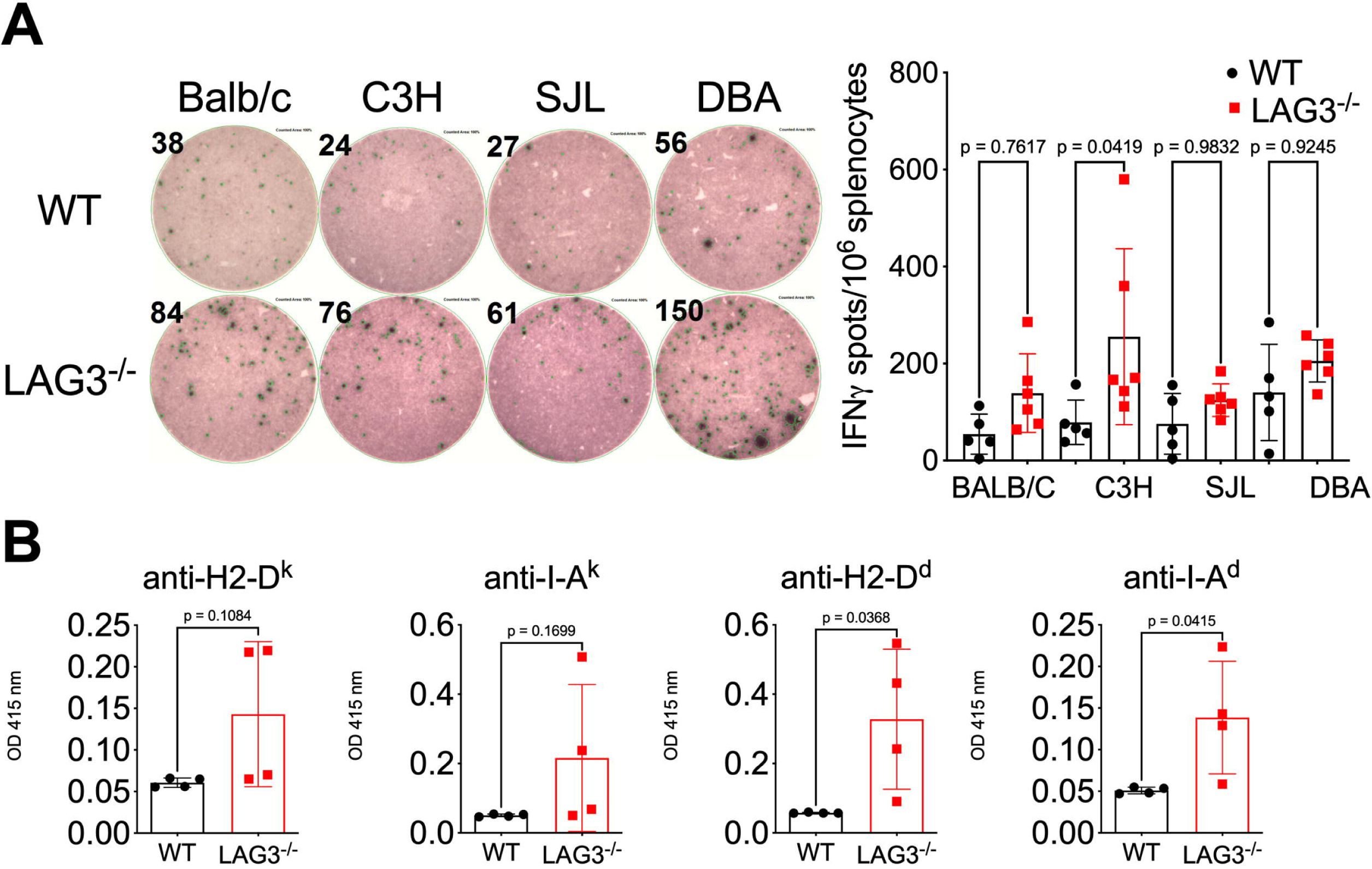
LAG3 deficient mice have increased alloreactivity. A. Left: Representative ELISPOT wells demonstrating the frequencies of alloreactive IFNγ-secreting splenic T cells in naïve non-transplanted B6.WT and B6.LAG3^-/-^ mice against BALB/c, C3H, SJL and DBA stimulator cells. Right: Quantification of basal frequencies of alloreactivity in B6.WT and B6.LAG3^-/-^ mice. **B.** ELISA quantification of serum IgG reactive to allogenic MHC I and II molecules. The data represents at least two pooled experiments where each symbol represents an individual mouse. Student’s T tests were performed and p<0.05 were considered significant.

### Recipient LAG3 deficiency enhances anti-donor alloimmune responses and elicits kidney allograft rejection

To test the functional consequences of LAG3 deletion, we used a previously described mouse model of kidney transplantation in which B6 (H-2^b^) recipients undergo bilateral nephrectomy, and are then transplanted with a single fully MHC-mismatched kidney allograft^43^. In this model the recipient survival is dependent upon allograft functionality, which can be measured by serum creatinine levels. Consistent with previously published studies that used different donor strains^43, 46^, B6.WT recipients spontaneously accept C3H renal allografts for >60 days (**Fig. 3A&B**). In contrast, most B6.LAG3^-/-^ recipients rapidly rejected kidney allografts (median survival time, MST, of 15d) (**Fig. 3B**) and had significantly increased serum creatinine levels at d. 14 (1.3 mg/dl versus 0.1 mg/dl in B6.WT controls and <0.4 mg/dl in non-transplanted mice, **Fig. 3C**). Immunohistochemistry of allografts recovered from LAG3^-/-^ recipients around the time of rejection (d. 14) showed decreased numbers of infiltrating T cells compared to grafts from B6.WT recipients (**Fig. S3**). Instead, the rejecting grafts showed characteristic signs of antibody mediated rejection including diffuse C4d staining, atrophic peritubular capillaries, endothelial cell swelling and edema (**Fig. 3D and Fig. S3**). Flow cytometry analysis of graft infiltrating cells on d. 10 posttransplant (**Fig. 3E-F**) revealed no significant changes in CD4^+^ or CD8^+^ T cell infiltration, with the only significant difference being in the increased numbers of infiltrating FoxP3^+^ T regulatory cells. LAG3^-/-^ recipients did demonstrate a modest increase in the infiltration of NK cells, the majority of which were of the NK1.1^hi^ phenotype (**Fig. 3E-F** and data not shown). This finding is consistent with antibody-mediated tissue injury of renal allografts as recently demonstrated by Yagisawa and colleagues^47^.

**Figure 3.**
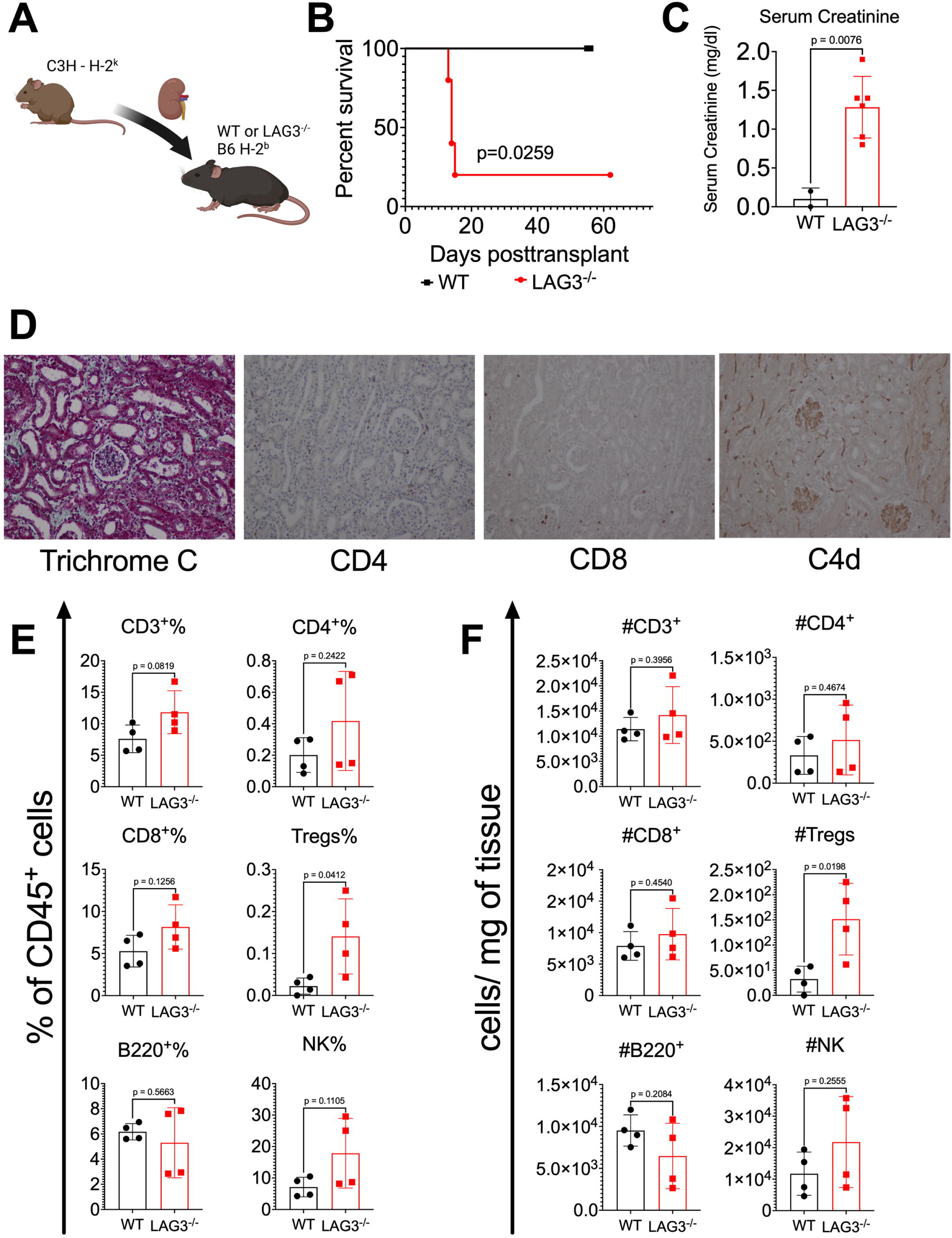
LAG3-deficient recipients acutely reject kidney allografts. Groups of B6.WT and B6.LAG3^-/-^ were transplanted with complete major histocompatibility complex-mismatched C3H kidney allografts (n=4-5/group). **A.** Renal allograft survival. **B.** Serum creatinine levels at d. 14 posttransplant. **C.** Renal allografts analyzed at the time of rejection (B6.LAG3^-/-^) or on d. 14 posttransplant (B6.WT) by Trichrome C and immunoperoxidase staining for CD4, CD8 and complement component C4d. The photographs were taken at 200x and are representative of 4-5 animals in each group. **D & E.** Flow cytometry analysis of graft infiltrating immune cells: CD3^+^ (CD45^+^ CD3^+^), CD4^+^ (CD45^+^ CD3^+^ CD4^+^), CD8^+^ (CD45^+^ CD3^+^ CD8^+^), Tregs (CD45^+^, CD3^+^ CD4^+^ FoxP3^+^), B220^+^ (CD45^+^ B220^+^), and NK (CD45^+^ CD3^-^ DX1^+^) at d10 following transplantation of C3H kidney allografts to B6.WT or B6.LAG3^-/-^ recipients. The data represents one of two experiments where each symbol represents an individual mouse. Statistical analysis of allograft survival was measured using Mantel-Cox log-rank test and for other analyses student’s T tests were performed and p<0.05 were considered significant.

The analysis of the peripheral T cell pool revealed no significant differences in the proportion or numbers of various spleen T cell subsets (**Fig. 4A** and **Fig. S4A**). Nevertheless, the absence of recipient LAG3 resulted in elevated frequencies of donor specific T cells, as measured by IFNγ ELIPSOT assay on d. 14 posttransplant (**Fig. 4C**). Analogous to the T cell compartment, spleen subsets of follicular, marginal zone, transitional, and regulatory B cells were similar in WT and LAG3^-/-^ allograft recipients (**Fig. 4B** and **Fig. S4B**), whereas germinal center B cells and plasma cells were increased in some but not all recipients. To assess anti-donor humoral immune responses, recipient serum was collected at 14 d posttransplant and analyzed for the levels of IgG antibodies against donor class I (D^k^) and class II (I-A^k^) MHC molecules. Consistent with our published studies, WT renal allograft recipients had developed minimal donor specific alloantbodies (DSA) at this timepoint. In contrast, LAG3^-/-^ recipients had significantly elevated levels of IgG against both D^k^ and I-A^k^ (**Fig. 4D**). Taken together these data demonstrate that LAG3 plays an essential role in regulating both cellular and humoral immune responses to transplanted allografts and contributes to spontaneous renal allografts acceptance in wild type recipients.

**Figure 4.**
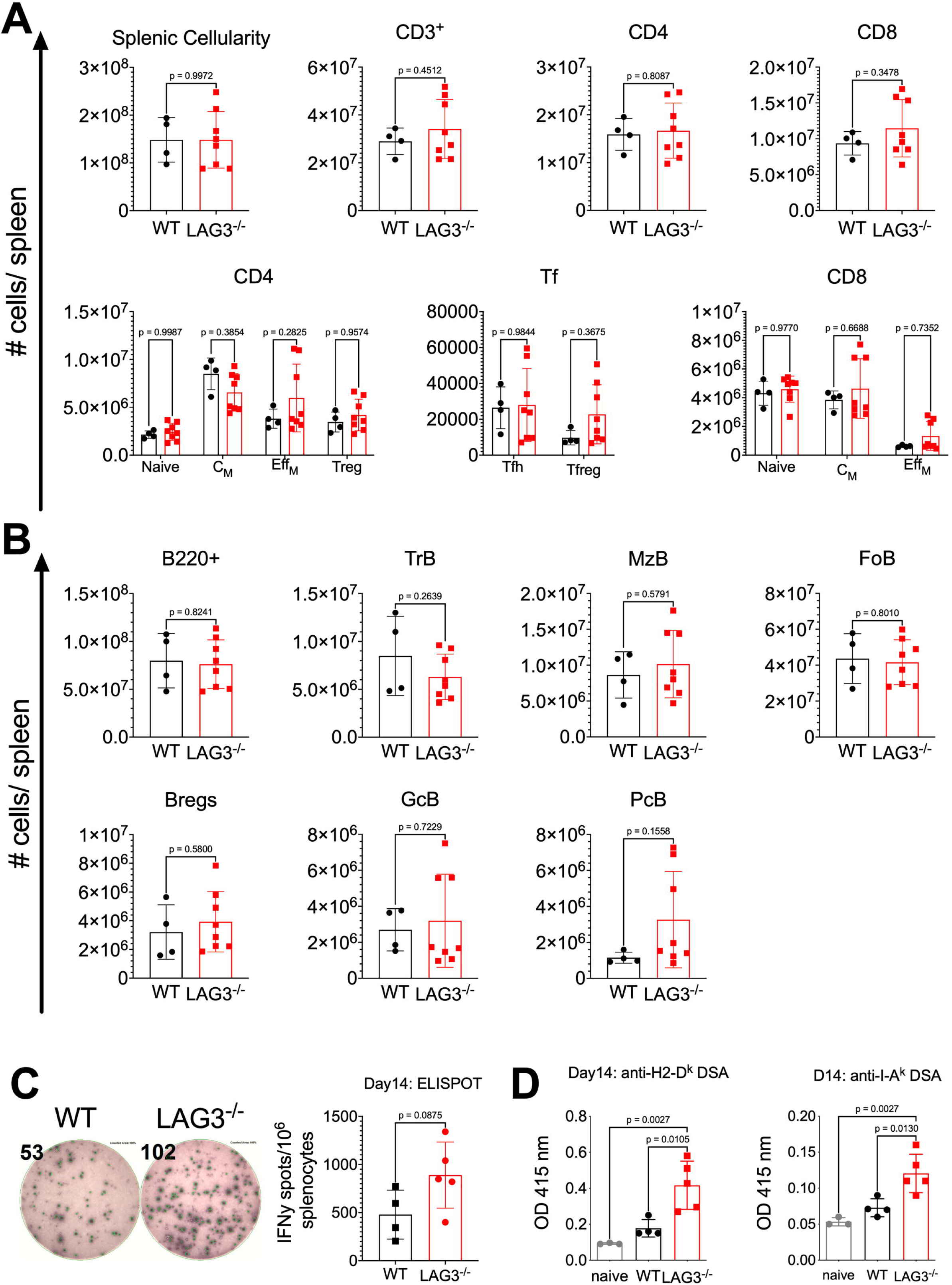
Recipient LAG3 deficiency enhances anti-donor immune responses. Analyses of donor-reactive immunity in B6.WT and B6.LAG3^-/-^ allograft recipients were performed at d. 10 posttransplant. **A.** The composition of spleen T cell subsets was defined as follows: CD3 – CD3^+^, CD4 – CD3^+^CD4^+^, CD8 – CD3^+^CD8^+^, Tregs – CD3^+^CD4^+^ FoxP3^+^, CD4 Naïve – CD3^+^CD4^+^CD62L^hi^CD44^lo^, CD4 C_M_ – CD3^+^CD4^+^CD62L^hi^CD44^hi^, CD4 Eff_M_ – CD3^+^CD4^+^CD62L^lo^CD44^hi^, CD8 Naïve – CD3^+^CD8^+^CD62L^hi^CD44^lo^, CD8 C_M_ – CD3^+^CD8^+^CD62L^hi^CD44^hi^, CD8 Eff_M_ – CD3^+^CD8^+^CD62L^lo^CD44^hi^, TFh – TCRb^+^CD4^+^FoxP3^-^PD-1^+^CXCR5^+^ and TFreg – TCRb^+^CD4^+^FoxP3^+^PD-1^+^CXCR5^+^. **B.** The composition of splenic B cell was defined as follows: B220 – B220^+^, FoB – B220^+^IgM^int^CD21/35^int^, MZB – B220^+^IgM^hi^CD21/35^hi^, TrB – B220^+^IgM^hi^CD21/35^lo^, Bregs – CD19^+^CD1d^hi^CD5^+^, GCB – B220^+^GL7^+^CD38^lo^, PCB – B220^-^CD138^hi^. **C.** The frequencies of donor reactive IFNγ-secreting splenocytes on d. 14 posttransplant. **D.** Serum levels of IgG against donor MHC-I (H2-D^k^) and MHC-II (I-A^k^). The data are pooled from two-three experiments, and each symbol represents an individual mouse. Student’s T tests were performed and p<0.05 were considered significant.

### B cells, but not CD8^+^T cells, are essential for kidney allograft rejection by LAG3^-/-^ recipients

Given that LAG3 is a well-studied regulator of T cell responses we anticipated that LAG3 deficiency primarily affects priming of alloreactive T cells and their pathogenic functions within the kidney graft. However, the modest increase in donor-reactive T cells and the graft histology (**Figs. 3-4 and Fig. S3)** suggested that alloantibody are the major mediators of graft tissue injury in LAG3^-/-^ recipients. To formally test the contributions of cellular vs antibody mediated rejection, LAG3^-/-^ recipients were treated with CD8^+^ T cell depleting antibody prior to transplantation of C3H kidney allografts (**Fig. 5A** and **Fig. S6B**). CD8^+^ T cell depletion antibodies failed to prolong kidney allograft survival in LAG3^-/-^ recipients, with MST of 16 days (**Fig. 5B**). Anti-donor humoral immune responses were assessed at 14 d posttransplant and analyzed for the levels of IgG antibodies against donor class I and class II MHC molecules. WT allograft recipients depleted of CD8 T cells had a low-grade DSA response at this timepoint. In contrast, LAG3^-/-^ recipients had significantly elevated levels of IgG against both donor class I and class II alloantigens (**Fig. 5C**). Although CD8 T cell depletion reduced the frequencies of IFNγ producing T cells in both groups, LAG3 deficient recipients still had modestly elevated frequencies of donor-specific T cells on d. 14 posttransplant compared to WT (**Fig. 5D**). This is likely due to a faster reconstitution of LAG3^-/-^ cells following depletion, which has been previously reported in memory CD4^+^ T cells^48^. Nevertheless, histological analysis at the time of rejection showed that grafts from CD8^+^ T cell depleted LAG3^-/-^ recipients had significant damage to the kidney tubules such as tubular dilation, casts, tubular atrophy, and edema. C4d staining of these grafts showed dilated capillaries, endothelial cell swelling, and heavy damage to the tubules and to the glomeruli (**Fig. 5E**). In summary, the grafts rejected by LAG3^-/-^ recipients following CD8^+^ T cell depletion displayed the typical features of antibody mediated graft damage similar to or even exceeding those observed in non-depleted LAG3-deficient recipients.

**Figure 5.**
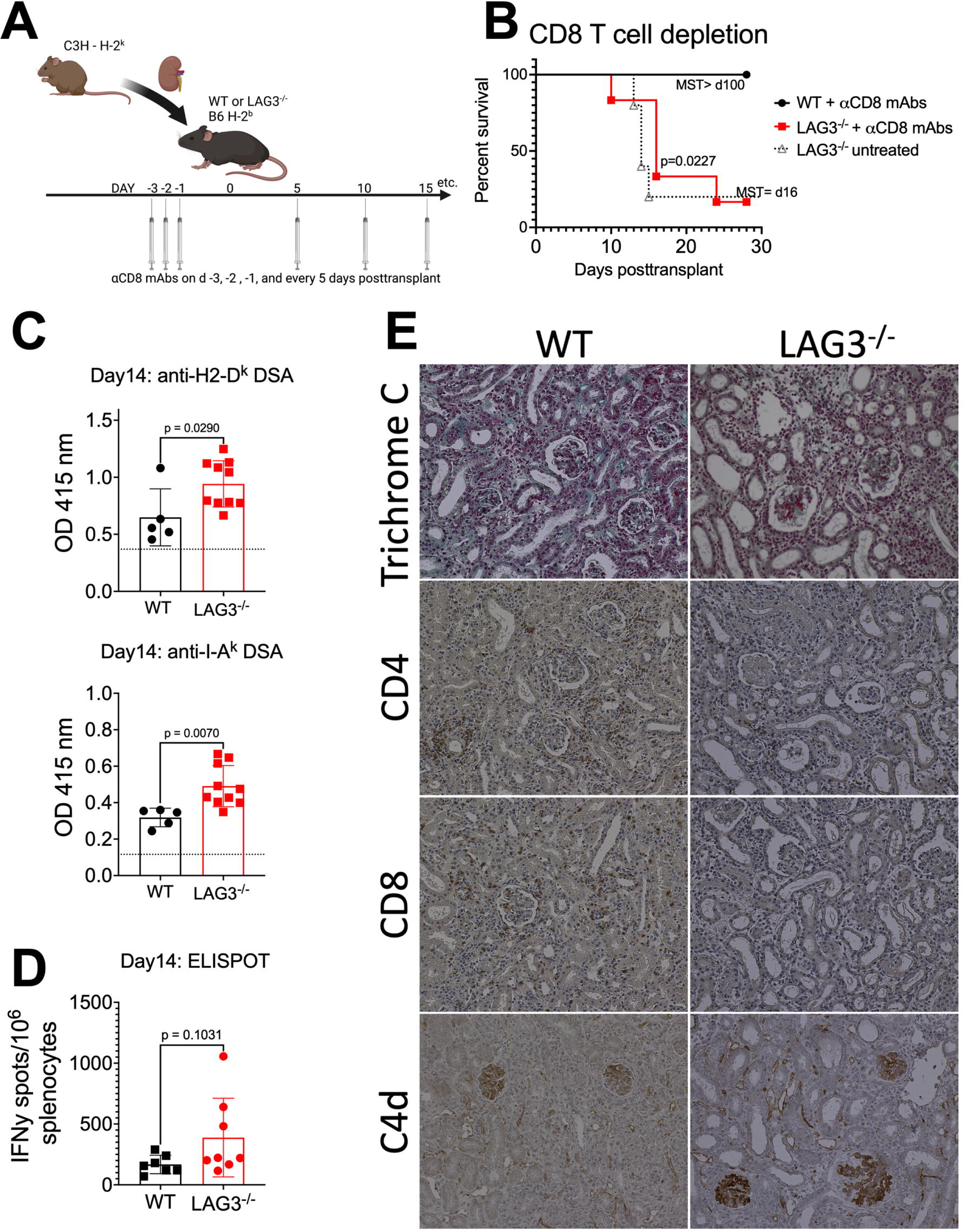
Kidney allograft rejection in LAG3 deficient recipients is not mediated by CD8 T cells. A. B6.WT or B6.LAG3^-/-^ recipients depleted of CD8^+^ T cells prior to transplantation of C3H renal allografts. **B.** Survival of renal allografts (n=5-6/group). **C.** Serum levels of IgG against donor MHC-I (H2-D^k^) and MHC-II (I-A^k^) in B6.WT and B6.LAG3^-/-^ kidney allograft recipients. **D.** The frequencies of donor reactive IFNγ-secreting splenocytes on d. 14 posttransplant. **E.** Renal allografts harvested at the time of rejection and analyzed by Trichrome C and immunoperoxidase staining for CD4, CD8 and complement component C4d. Images were taken at 200x and are representative of 4-5 animals in each group. The data are pooled from two-three experiments, and each symbol represents an individual mouse. Statistical analysis of allograft survival was measured using Mantel-Cox log-rank test and for other analyses student’s T tests were performed and p<0.05 were considered significant.

To test the roles of B cells and antibodies in the observed rejection, LAG3^-/-^ recipients of C3H kidney allografts were treated with anti-mouse CD19 and B220 B cell depleting antibodies starting on d. 3 posttransplant and throughout the experiment (**Fig. 6A** and **Fig. S6A**). Remarkably, B cell depletion restored graft survival in the majority of LAG3^-/-^ recipients (**Fig. 6B**). Depletion of recipient B cells lead to the abrogation of the DSA responses in both WT and LAG3^-/-^ recipients (**Fig. 6C**). While B cell depletion reduced T cell alloresponses in both groups, LAG3^-/-^ recipients still had increased frequencies of donor-reactive IFNγ producing T cells compared to WT (**Figs. 6D and 4C**). Histological analysis at d. 30 posttransplant showed that recipient B cell depletion reduced allograft damage with marked decrease in T cell infiltrates and antibody binding, demonstrated by the absence of C4d staining (**Fig. 6E**). Together, these results demonstrate that the rejection observed in LAG3^-/-^ mice is driven by B cell production of DSA and not by cytotoxic T cells, and that the generated DSA is a major effector mechanism of graft injury.

**Figure 6.**
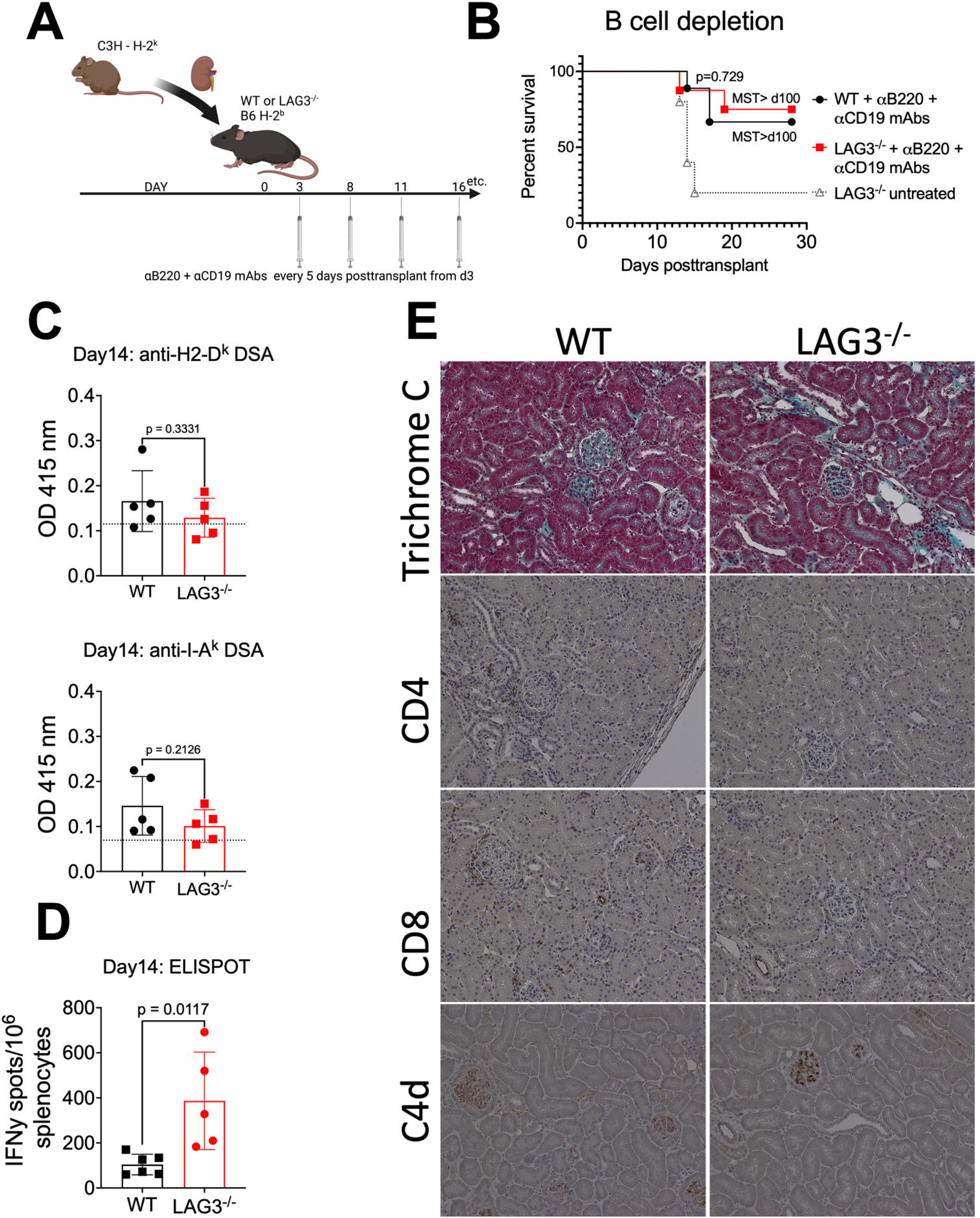
Kidney allograft rejection by LAG3 deficient recipients is mediated by B cells. A. B cells were depleted in B6.WT or B6.LAG3^-/-^ recipients after transplantation of C3H renal allografts. **B.** Survival of renal allografts (n=5-6/group). **C.** Serum levels of IgG against donor MHC-I (H2-D^k^) and MHC-II (I-A^k^). **D.** The frequencies of donor reactive IFNγ-secreting splenocytes on d. 14 posttransplant. **E.** Renal allografts harvested at the time of rejection and analyzed by Trichrome C and immunoperoxidase staining for CD4, CD8 and complement component C4d. Images were taken at 200x and are representative of 4-5 animals in each group. The data are pooled from two-three experiments, and each symbol represents an individual mouse. Statistical analysis of allograft survival was measured using Mantel-Cox log-rank test and for other analyses student’s T tests were performed and p<0.05 were considered significant.

### LAG3 is expressed by regulatory B cells and plasma cells in kidney allograft recipients

As our data revealed that B cell responses are affected by the loss of LAG3 mediated immune regulation, we next analyzed LAG3 expression in different B cell subsets following transplantation. LAG3 expression was not detectable by flow cytometry in B cells from naïve mice (data not shown) consistent with previous reports that LAG3 expression on B cells is only inducible in a T cell dependent manner^3^. Given the observed rejection kinetics (**Fig. 3B & 5B**), we measured LAG3 expression on splenic B cell subsets on d10 posttransplant, using LAG3^-/-^ recipients as a control. Of the cell populations tested, only 16.28% (±5.227 SD) CD19^+^CD1d^hi^CD5^+^ Bregs and 8.08% (±1.979 SD) CD138^+^ plasma cells expressed cell surface LAG3 (**Fig. 7A & B** and **Fig. S5**), suggesting differential expression in specialized subsets.

**Figure 7.**
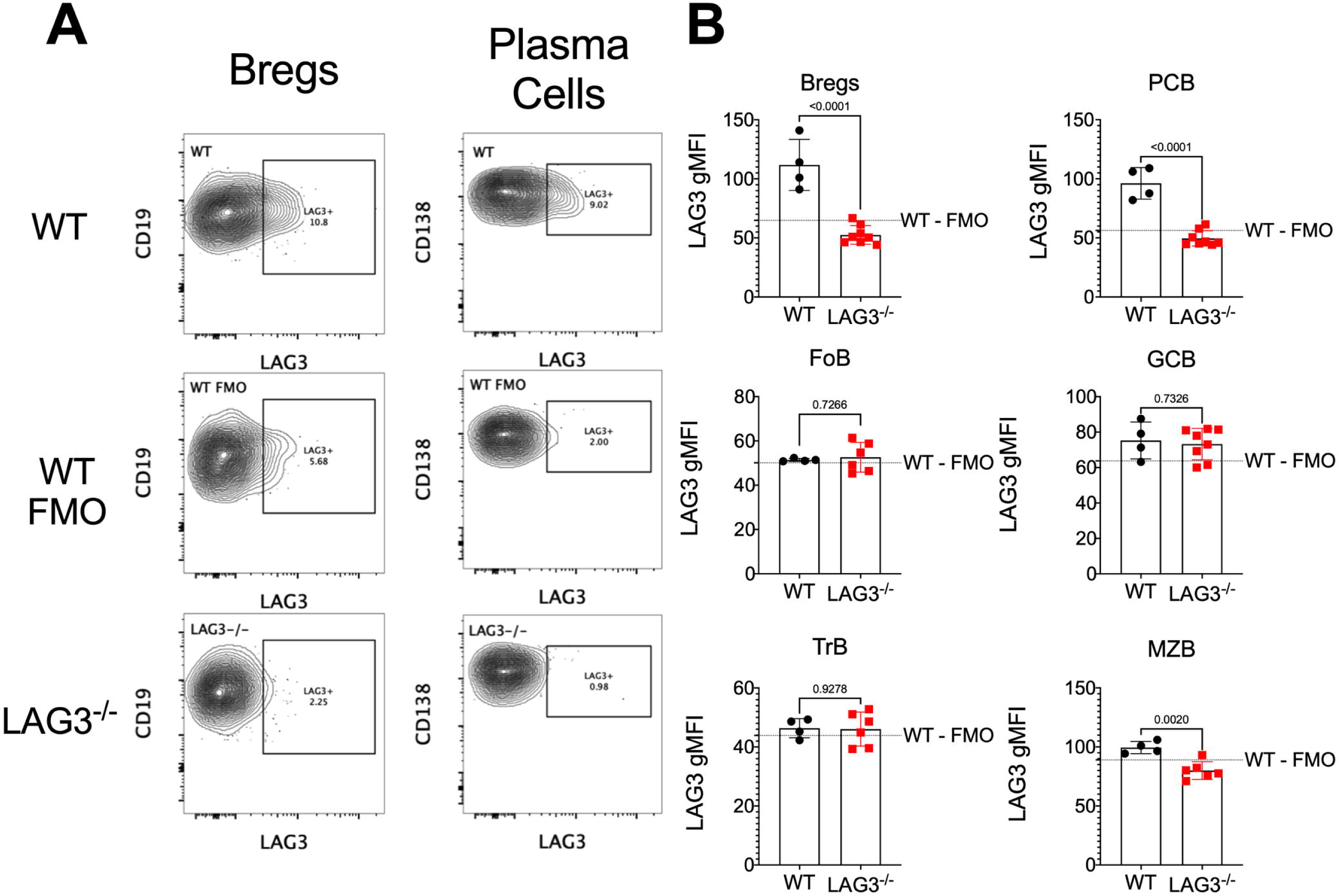
LAG3 is induced on B cell subsets following transplantation. Analyses of LAG3 expression on B cell subsets in B6.WT and B6.LAG3^-/-^ renal allograft recipients on d. 10 posttransplant. **A.** Representative contour plots of LAG3 expression in Bregs and plasma cell pool. **B.** Quantification of LAG3 expression in different B cell subsets following transplantation. The dashed line represents the mean MFI of the WT-FMO control. The data are pooled from two experiments, and each symbol represents an individual mouse. Student’s T tests were performed and p<0.05 were considered significant.

### LAG3 deficiency on both T and B lymphocytes is required for kidney allograft rejection

The absence of LAG3 on T cells often induces pathogenesis in other mouse models ^15^. To address whether increased T cell help or defective T regulatory cell function was a major driver of the observed allograft injury, we used T cell conditional knockout mice and littermate controls as renal allograft recipients (**Fig. 8A**). B6.CD4Cre^+/-^LAG3^fl/fl^ which lack LAG3 on CD4^+^ and CD8^+^ T cells spontaneously accepted kidney allografts for > 60 days (**Fig. 8B**) indicating that LAG3 deficiency on T cells alone was not sufficient to induce rejection. Analysis of the immune response in these recipients showed elevated DSA responses (**Figs. 8C**), but no significant changes in the frequencies of IFNγ producing alloreactive T cells (**Fig. 8D**). Histological analysis of the allografts at four weeks posttransplant revealed no major impact on fibrosis or graft tissue injury Interestingly, despite the presence of serum DSA, T cell LAG3 deficiency resulted in reduced C4d deposition within interstitial capillaries compared to LAG3^-/-^ recipients (**Figs. 8E and 3D**).

**Figure 8.**
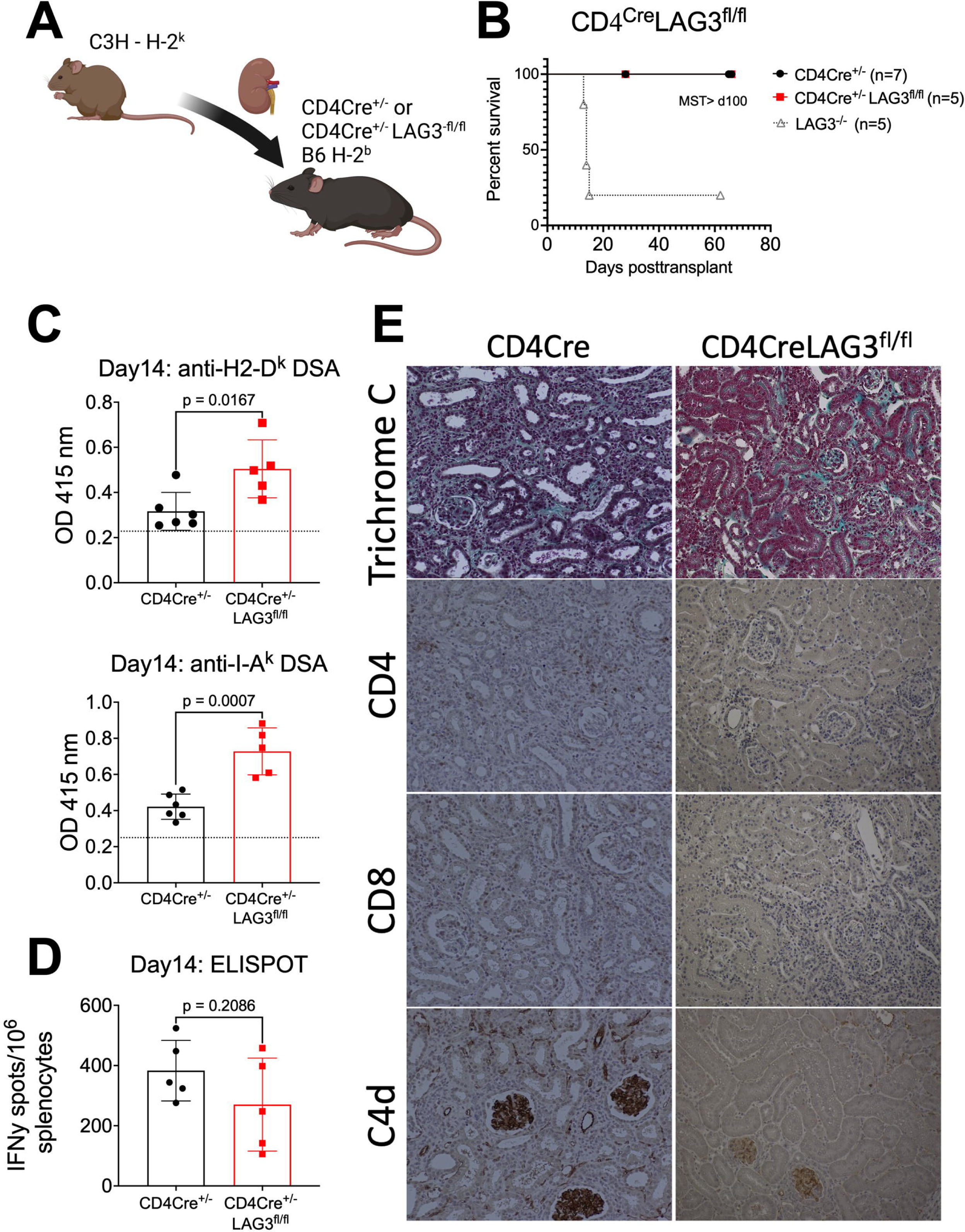
T cell LAG3 expression is dispensable for kidney allograft acceptance. A. Experimental design. **B.** Survival of renal allografts (n=5-7/group). **C.** Serum levels of IgG against donor MHC-I (H2-D^k^) and MHC-II (I-A^k^) in B6.CD4Cre^+/-^LAG3^fl/fl^ or B6.CD4Cre^+/-^ littermate control kidney allograft recipients. **D.** The frequencies of donor reactive IFNγ-secreting splenocytes on d. 14 posttransplant. **E.** Renal allografts harvested at the time of rejection and analyzed by Trichrome C and immunoperoxidase staining for CD4, CD8 and complement component C4d. Images were taken at 200x and are representative of 4-5 animals in each group. The data are pooled from two-three experiments, and each symbol represents an individual mouse. Statistical analysis of allograft survival was measured using Mantel-Cox log-rank test and for other analyses student’s T tests were performed and p<0.05 were considered significant.

We next investigated the role of LAG3 using a B cell conditional knockout recipient model (**Fig. 9A**). Unexpectedly, B6.CD19Cre^+/-^LAG3^fl/fl^ recipients with specific LAG3 deficiency in B lymphocytes also failed to reject kidney allografts by d. 30 posttransplant (**Fig. 9B**). LAG3 deficiency on B cells of renal transplant recipients resulted in a modest increase in DSA levels by d14 posttransplant (**Fig. 9C**). In contrast, LAG3 deficiency on B cells had no impact on the frequencies of donor reactive T cells at d14 posttransplant (**Fig. 9D**), indicating that B cell LAG3 expression does not affect T cell priming. Graft histology at four weeks posttransplant showed no significant differences in T cell infiltration. C4d staining was less intense in the CD19Cre^+/-^ LAG3^fl/fl^ recipients compared to the littermate controls but showed a rarefaction of peritubular capillaries, and endothelial cell swelling consistent with antibody mediated graft injury (**Fig. 9E** and **Fig. S7**)^49^. Taken together these data show that LAG3 expression on either helper T cells or B cells is sufficient to induce spontaneous acceptance of murine kidney allografts.

**Figure 9.**
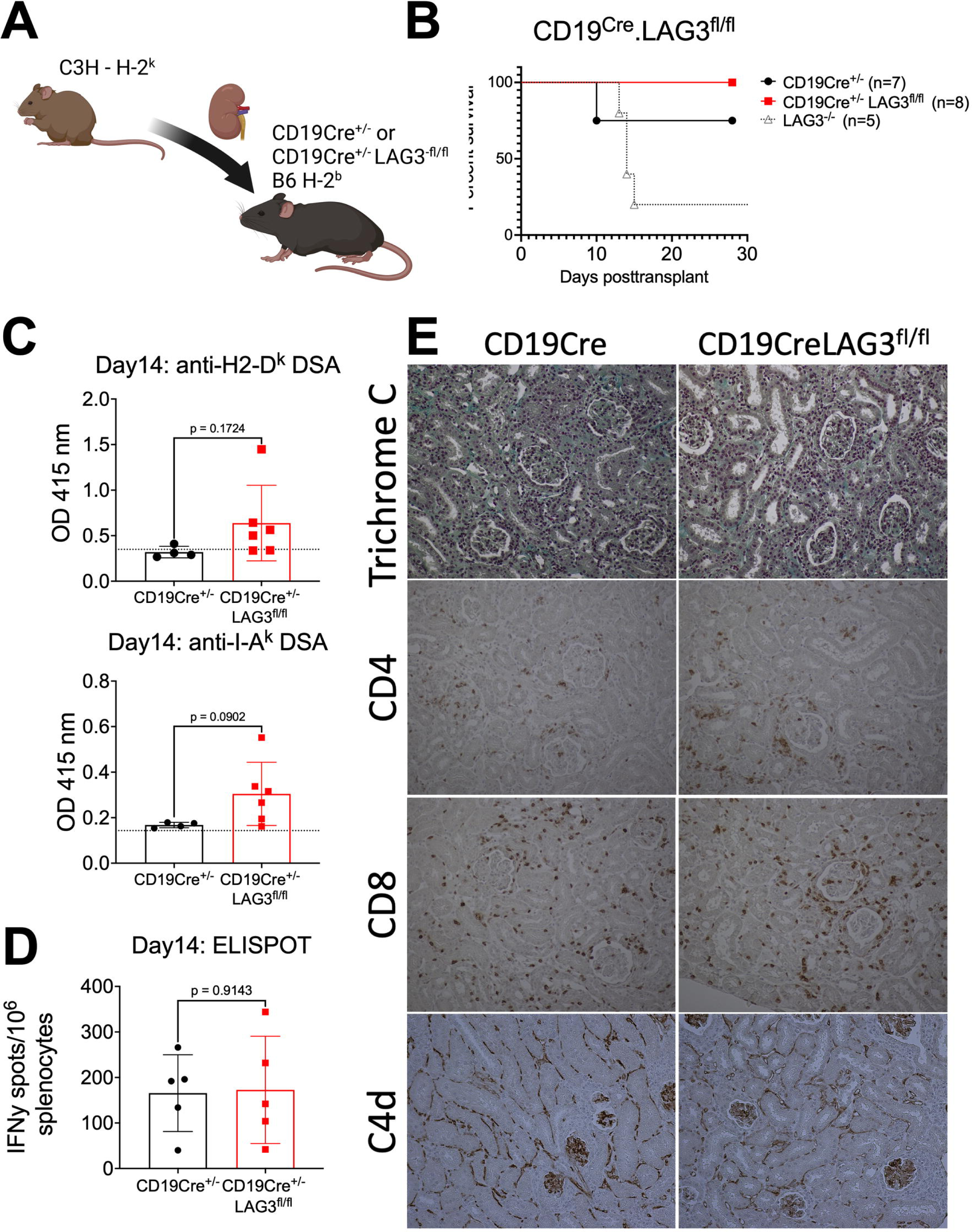
B cell LAG3 expression is dispensable for kidney allograft acceptance. A. Experimental design. **B.** Survival of renal allografts (n=7-8/group). **C.** Serum levels of IgG against donor MHC-I (H2-D^k^) and MHC-II (I-A^k^) in B6.CD19Cre^+/-^ littermate controls or B6.CD19Cre^+/-^LAG3^fl/fl^ kidney allograft recipients. **D.** The frequencies of donor reactive IFNγ-secreting splenocytes on d. 14 posttransplant. **E.** Renal allografts harvested at the time of rejection and analyzed by Trichrome C and immunoperoxidase staining for CD4, CD8 and complement component C4d. Images were taken at 200x and are representative of 4-5 animals in each group. The data are pooled from two-three experiments, and each symbol represents an individual mouse. Statistical analysis of allograft survival was measured using Mantel-Cox log-rank test and for other analyses student’s T tests were performed and p<0.05 were considered significant.

### LAG3 regulates de novo alloresponses

Our data show that non-transplanted LAG3 deficient mice have increased alloreactive immune responses (**Fig. 2A&B**). To establish whether LAG3 is regulating *de novo* immune responses, we treated WT kidney allograft recipients with a course of LAG3 blocking antibody and monitored mice for signs of rejection (**Fig. 10A**). Blockade of LAG3 resulted in rejection in 2 out 9 mice within 14 days posttransplant (**Fig. 10B**). Despite the long term graft survival in the remaining recipients, we found that LAG3 blockade did induce graft damage, as measured by kidney injury markers NGAL and KIM1 in the urine, and blood urea nitrogen (BUN) in the serum (**Fig. 10C-E**). Blockade of LAG3 also resulted in increased DSA production (**Fig. 10F&G**) and increased frequencies of donor reactive T cells (**Fig. 10H**), indicating that LAG3 regulates de novo alloresponses following transplantation. The increase in markers of kidney injury and alloresponses suggested that LAG3 blockade may induce chronic graft injury. Indeed, histological analysis at d. 42 posttransplant (**Fig. 10I & Fig. S8**) revealed tubular atrophy, endothelial cell swelling and increased alpha-smooth muscle actin (aSMA) staining due to graft fibrosis, also reflected in the Trichrome staining (**Fig. S8**). C4d staining showed C4d deposition in both control IgG treated and anti-LAG3 treated recipients, but recipients that received LAG3 blockade had dilated peritubular capillaries with marginated mononuclear cells (red arrows, **Fig. 10I**). In addition, recipients that received LAG3 blockade had increased CD4^+^ and CD8^+^ T cell graft infiltrates. These findings identify LAG3 as an important regulator of *de novo* immune responses to a kidney allograft.

**Figure 10.**
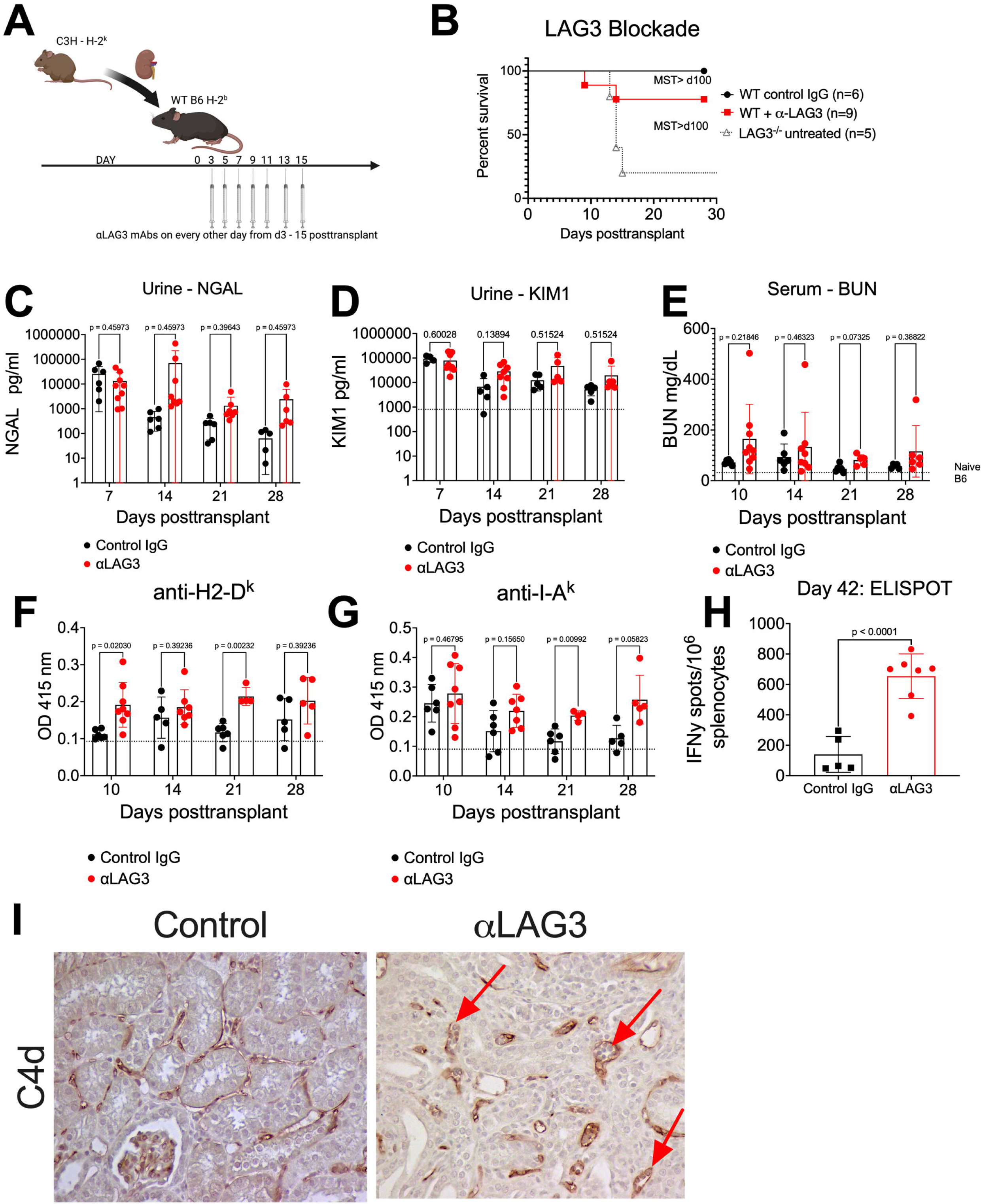
**LAG3 blockade enhances de novo alloresponses following kidney transplantation leading to chronic antibody mediated graft injury**. **A**. B6.WT treated with anti-LAG3 mAb (Clone C9B7W) or control IgG posttransplantation of C3H renal allografts. **B**. Survival of renal allografts (n=6-9/group). **C-D**. NGAL and KIM1 levels in the urine collected from kidney allograft recipients. **E**. Serum Blood Urea Nitrogen (BUN) levels. **F-G**. Serum levels of IgG against donor MHC-I (H2-D^k^) and MHC-II (I-A^k^). **H**. The frequencies of donor reactive IFNγ-secreting splenocytes on d. 42 posttransplant. **I**. Renal allografts harvested at the time of rejection and analyzed by immunoperoxidase staining for complement component C4d. Images were taken at 400x and are representative of 4-5 animals in each group. The data are pooled from two-three experiments, and each symbol represents an individual mouse. Statistical analysis of allograft survival was measured using Mantel-Cox log-rank test, for the time-course analysis of kidney injury markers and for DSA one-way ANOVA with Tukey’s multiple comparison was performed and for ELISPOT analysis student’s T tests were performed and p<0.05 were considered significant.

## Discussion

Coinhibitory molecule LAG3 is best studied in regulating effector and regulatory T cell functions in autoimmunity, infection and cancer^17–22^. However, little is known about the contribution of this pathway in regulating humoral immune responses. Our results definitively demonstrate that LAG3 regulates alloantibody generation in response to solid organ transplantation, and suggest that LAG3 expression on both T and B lymphocytes plays a role in this process.

The absence of LAG3 in T cells leads to enhanced effector T cell responses and increased formation of T cell memory^26^. Consistent with this, naïve, non-transplanted LAG3-deficient mice have elevated levels of CD44^hi^ memory T cells and enhanced response to a panel of alloantigens compared to wild type counterparts (**Fig. 1 & Fig. 2**). In addition, we observed increased numbers of germinal center B cells and CD138^+^ plasma cells in naïve 10 week old LAG3^-/-^ mice, which suggested ongoing B cell activation. Based on previous studies, we initially proposed that LAG3 deletion in the recipient will result in dysregulated alloimmunity characterized by exaggerated activation of alloreactive T cells and leading to T cell-mediated rejection.

The model of renal transplantation was chosen for these experiments as even fully MHC mismatched mouse kidney allograft typically do not undergo acute rejection, and are spontaneously accepted in many donor/recipients strain combinations^50^. While functional C3H kidney allografts survive for > 60 days in B6.WT recipients, they are rapidly rejected by B6.LAG3^-/-^ mice (**Fig. 2**). Long term survival and function of mouse renal allografts have been previously correlated with the numbers of graft-infiltrating regulatory T cells, as FoxP3^+^ regulatory T cell depletion results in rapid graft rejection^51, 52^. In contrast to germline LAG3^-/-^ mice, T cell specific LAG3 deletion did not result in acute graft rejection (**Fig. 8**). LAG3 is highly expressed by FoxP3^+^ Tregs and LAG3 is thought to play a role in regulatory T cell’s optimal suppressor activity^6, 53, 54^. Previous studies demonstrated that FoxP3 regulatory T cell depletion in mouse renal allograft recipients resulted in T cell mediated rejection without affecting DSA generation^52^. Moreover, LAG3^-/-^ mice have increased numbers of FoxP3^+^ T cells before and after transplantation (**Figs. 1-3**). However, it is unlikely that the augmented alloimmunity and graft rejection in our model are entirely due to dysfunctional Tregs as specific deletion of LAG3 in recipient T cells (including Tregs) was not sufficient to induce rejection (**Fig. 8**). ^52^.

While LAG3 expression in B cells was reported in 2005^3^, its contribution to their functions has not been extensively studied. Our results demonstrate that the lack of recipient LAG3 leads to increased titers of IgG DSA antibodies and allograft rejection with characteristic features of antibody-mediated injury. Furthermore, depletion experiments confirmed that B cells are required for acute rejection of kidney allografts (**Fig. 6**), whereas the rejection still occurred with the same kinetics in CD8 T cells-depleted LAG3^-/-^ recipients (**Fig. 5**). To date, there are only a few studies addressing the role of LAG3 in humoral immune response. Butler and colleagues^55^, reported that the combination treatment with anti-PD-L1 and anti-LAG3 mAb during established malaria enhances Tfh responses, plasma cell formation and protective antibody generation in a mouse model of malaria. However, the individual contributions of PD-L1 and LAG3 blockade were not evaluated in this study. Another recent study identified a novel subset of LAG3^+^ IL-10 secreting plasma cells with regulatory properties has recently been reported^27^. However, the relative importance of LAG3 on helper T cells vs B cells during the initiation of B cell activation, germinal center formation and differentiation into antibody-secreting cells remains to be determined and is the focus of ongoing studies in our laboratory.

Our study does not entirely rule out the contribution of LAG3 expressed by cells other than T and B lymphocytes. NK cells express LAG3 and are important mediators of antibody-mediated injury of renal allografts^47^. While the effects of LAG3 on various NK cell functions is highly controversial (reviewed in^56^), it is possible that dysregulated NK cells mediate rejection in LAG3^-/-^ recipients, secondary to enhanced T cell activation and DSA generation. The absence of LAG3 on recipient antigen presenting cells (APCs) may also contribute to elevated T cell responses. Initial analyses of LAG3 expression in dendritic cell (DC) subsets showed high LAG3 expression by plasmacytoid but not lymphoid or myeloid conventional DCs^7^. However, a more recent study demonstrated a role for LAG3 on bone marrow derived DCs in optimal T cell priming^57^. Due to these controversies, the impact of LAG3 deficiency in recipient APCs needs to be carefully dissected in future studies.

It is important to note that, treatment of WT B6 recipients with a LAG3 blocking antibody induced a pathology consistent with chronic rejection (**Fig. 10**). This was evident in the increased kidney injury markers in the urine and serum (**Fig. 10C-E**), increased serum DSA levels (**Fig.10F&G**), and increased histological signs of injury (**Fig. 10I** and **Fig. S8**). There are a paucity of small animal chronic injury/rejection models in kidney transplantation, with some of the current models relying on repeated administration of anti-MHC-I antibodies^58^, or repeat transfers of anti-donor antibody containing sera^59, 60^. Our findings thus identify a new clinically relevant model for studying the mechanisms of chronic injury and graft rejection in renal transplantation.

To our knowledge, our study is one of the first to address the role of LAG3 during alloimmune responses. Lucas *et al*. reported that blocking LAG3 with mAb prevents tolerization of CD8 T cells following allogeneic bone marrow transplantation in mice^38^. In an earlier study, the investigators used depleting anti-LAG3 mAb as an induction therapy in a rat model of cardiac transplantation. Interestingly, the treatment extended heart allograft survival, yet abrogated the tolerogenic effects of donor specific cell transfusion, which the authors attributed to Treg depletion^39^.

Despite many gaps and controversies in LAG3 biology, it is a molecule of therapeutic interest, particularly in the cancer field. Antagonistic antibodies such as Relatlimab are given to patients in combination with anti-PD-1 therapy in phase II clinical trials^61, 62^. Another promising reagent undergoing phase III trials is a bispecific antibody against LAG3 and PD-1, MGD013^28^. Anti-LAG3 depleting antibodies have been developed and validated in non-human primates to target overactivated immune cells during undesired immune response^63^. An agonistic anti-LAG3 antibody (IMP761) is currently under development for T cell mediated autoimmunity, and this approach could have potential for clinical transplantation^20^.

In conclusion, our study demonstrates for the first time the importance of LAG3 in regulating pathogenic humoral immune responses to a transplanted solid organ. The results provide rationale for investigating LAG3 impact on B cell activation, survival and differentiation, and suggest LAG3 as a potential target for future therapeutic interventions for the prevention and treatment of antibody-mediated rejection.

## Author Contributions

Design of research study: MN, JL, WMB, EC, MA, RLF, BM & AV

Conducting Experiments: MN, RF, JL, VG, JIV, AB, & ND

Acquiring data: MN, JL, VG, JIV, & AB

Analyzing data: MN, JL, VG, JIV, & WMB

Providing reagents/mice: BM

Writing the manuscript: MN, WB, BM, & AV

## Acknowledgements

Models were created with Biorender.com. This work was supported by funding from the American Society of Transplantation (gCDX-221C0MN – MN), and the National Institutes of Health (R01AI165513 – WMB, R01AI-74740 – RLF, R01AI125247 – BM, R01AI60740 & R01AI113142-AV).

