## Supplementary Figures for "LAG3 regulates antibody responses in a murine model of kidney transplantation"

Figure S1.

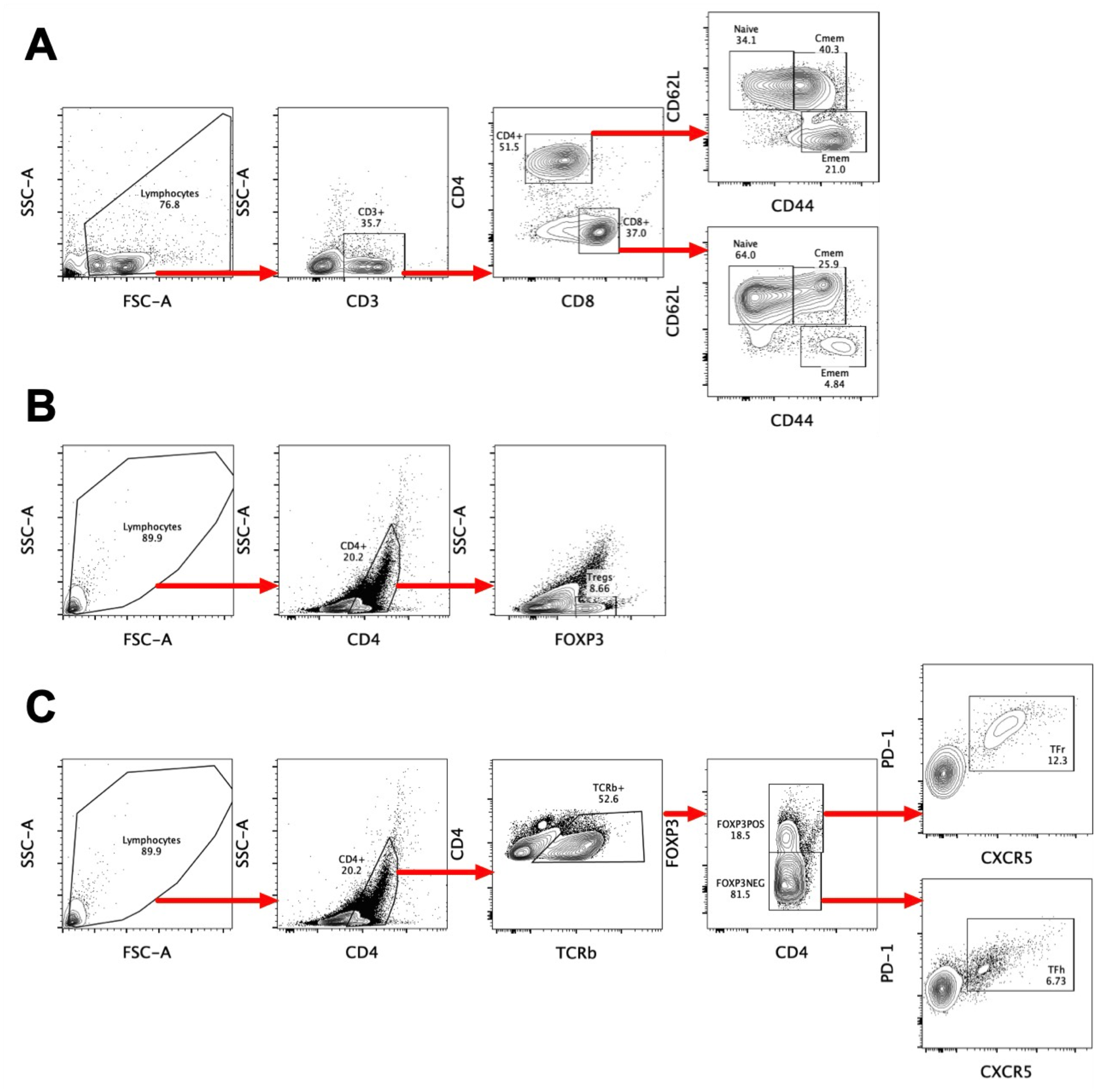

**Figure S1. Splenic T cell gating strategies**. ***A****. CD4 (CD3+CD4+), CD8 (CD3+CD8+), CD4 TEffM (CD3+, CD4+, CD62Llo, CD44hi), CD8 TEffM (CD3+, CD8+, CD62Llo, CD44hi).* ***B****. Tregs (CD3+, CD4+, FoxP3+).* ***C****. Tfh (TCRb+, CD4+, FoxP3-, PD-1+, CXCR5+) and Tfr cells (TCRb+, CD4+, FoxP3+, PD-1+, CXCR5+).*

Figure S2.

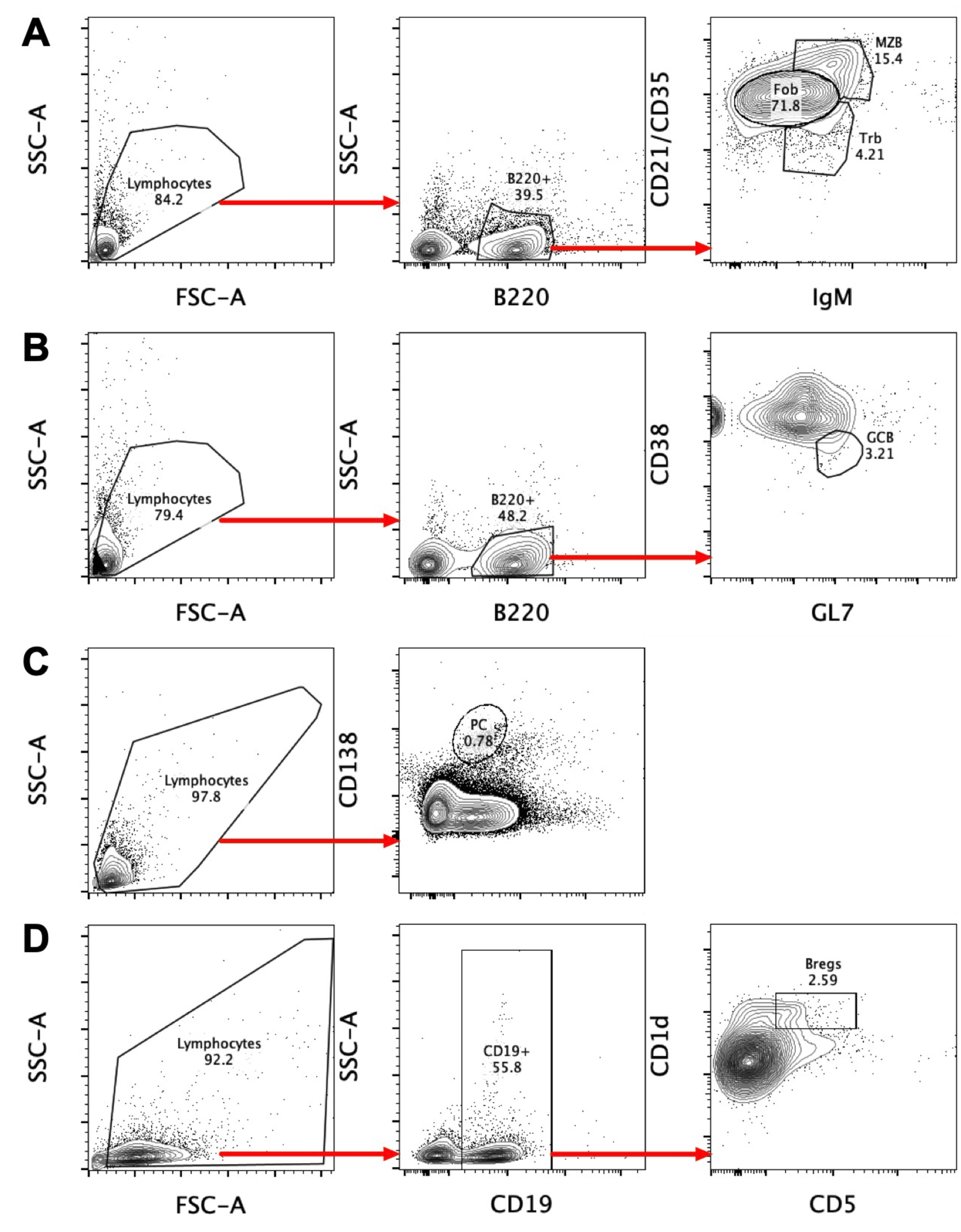

***Figure S2. Splenic B cell gating strategies****.* *Subsets were defined as follows;* ***A****. FoB* -*B220^+^, IgM^int^, CD21/35^int^, MZB - B220^+^, IgM^hi^, CD21/35^hi^,TrB - B220^+^, IgM^hi^, CD21/35^lo^.* ***B****. GCB - B220^+^, GL7^+^, CD38^lo^.* ***C****. PCB –B220^-^, CD138^hi^.* ***D****. Bregs - CD19^+^,CD1d^hi^, CD5^+^.*

Figure S3

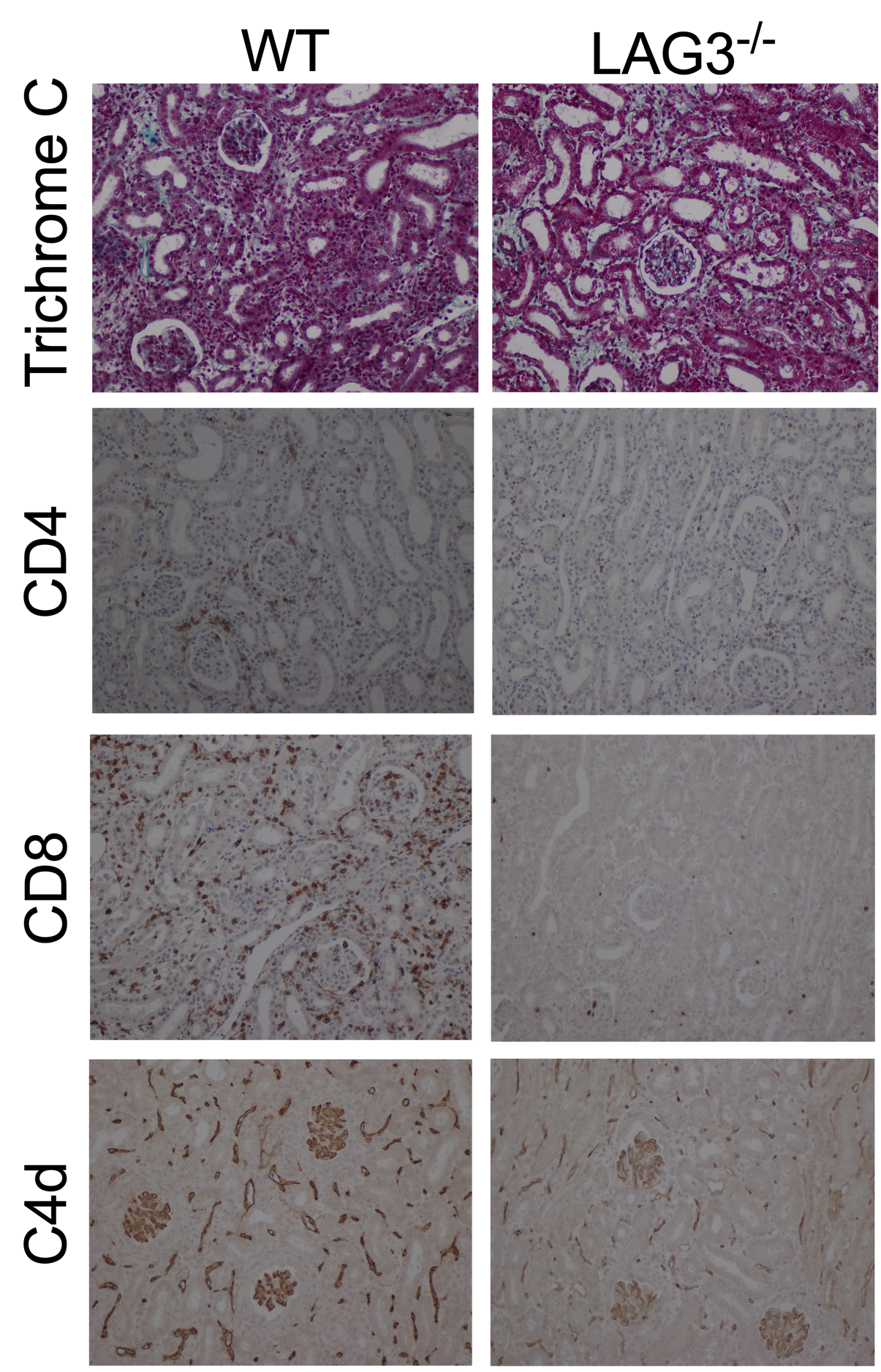

***Figure S3. Graft histology of WT and LAG3-/- renal allogratfts****. Renal allografts analyzed at the time of rejection (B6.LAG3^-/-^) or on d. 14 posttransplant (B6.WT) by Trichrome C and immunoperoxidase staining for CD4, CD8 and complement component C4d. The photographs were taken at 200x and are representative of 4-5 animals in each group.*

Figure S4

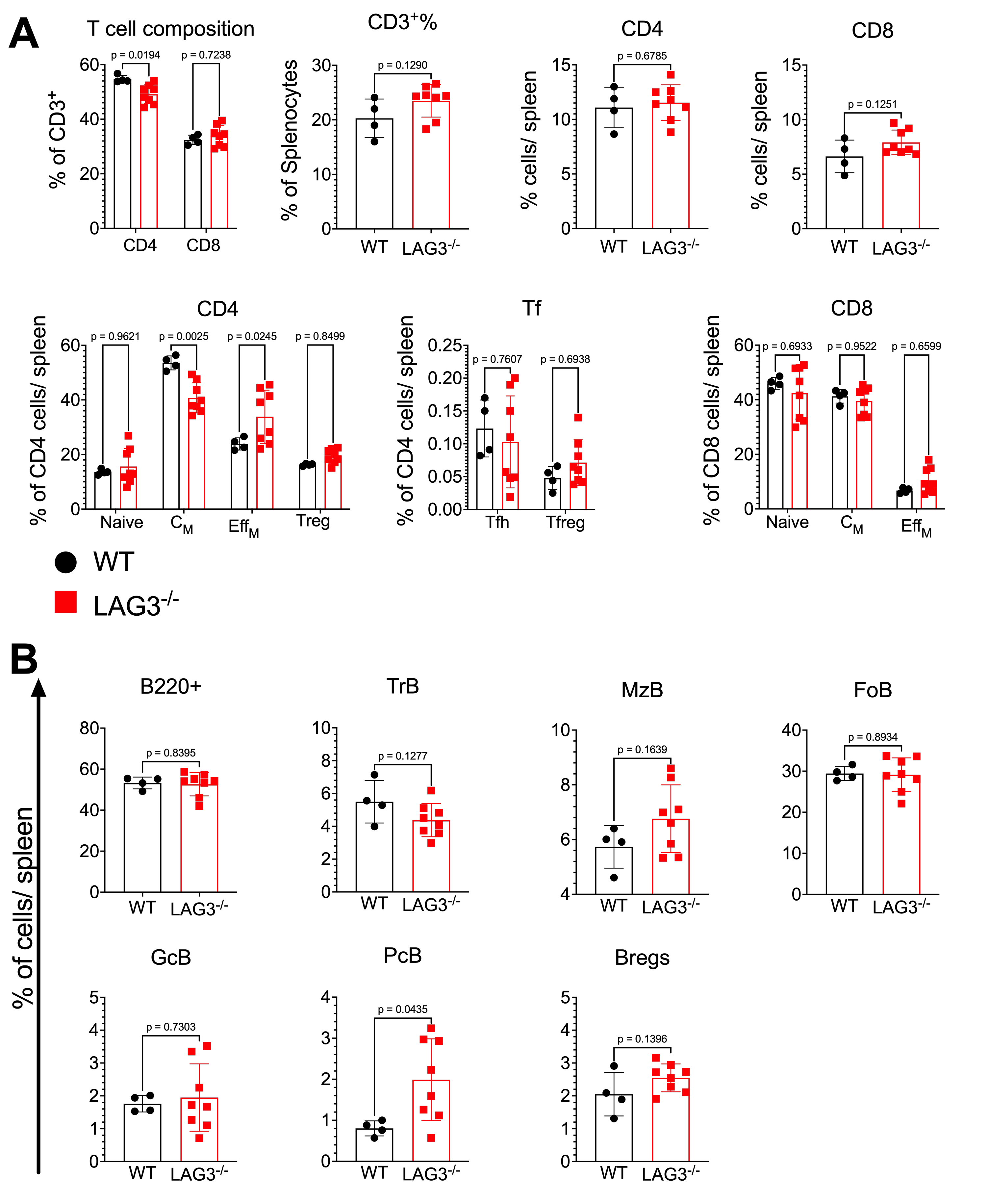

***Figure S4. Recipient LAG3 deficiency enhances anti-donor immune responses.*** *Analyses of donor-reactive immunity in B6.WT and B6.LAG3^-/-^ allograft recipients were performed at d. 10 posttranplant.* ***A.*** *The frequencies of spleen T cell subsets defined as follows: CD3 – CD3^+^, CD4 – CD3^+^CD4^+^, CD8 - CD3^+^CD8^+^, Tregs - CD3^+^CD4^+^ FoxP3^+^, CD4 Naïve - CD3^+^CD4^+^CD62L^hi^CD44^lo^, CD4 C_M_ - CD3^+^CD4^+^CD62L^hi^CD44^hi^, CD4 Eff_M_ - CD3^+^CD4^+^CD62L^lo^CD44^hi^, CD8 Naïve - CD3^+^CD8^+^CD62L^hi^CD44^lo^, CD8 C_M_ - CD3^+^CD8^+^CD62L^hi^CD44^hi^, CD8 Eff_M_ - CD3^+^CD8^+^CD62L^lo^CD44^hi^, TFh - TCRb^+^CD4^+^FoxP3^-^PD-1^+^CXCR5^+^ and TFreg - TCRb^+^CD4^+^FoxP3^+^PD-1^+^CXCR5^+^.* ***B.*** *The composition of splenic B cell defined as follows: B220 – B220^+^, FoB* - *B220^+^IgM^int^CD21/35^int^, MZB - B220^+^IgM^hi^CD21/35^hi^, TrB - B220^+^IgM^hi^CD21/35^lo^, Bregs - CD19^+^CD1d^hi^CD5^+^, GCB - B220^+^GL7^+^CD38^lo^, PCB - B220^-^ CD138^hi^. Student’s T tests were performed and p<0.05 were considered significant.*

Figure S5

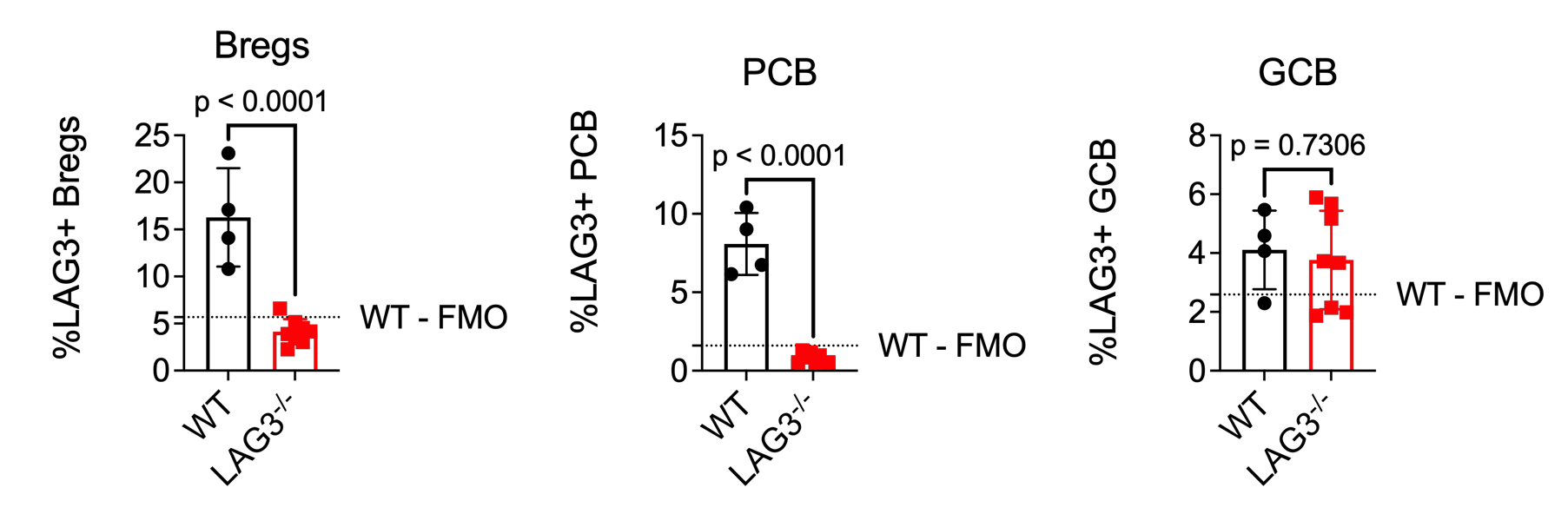

***Figure S5. LAG3 is induced on B cell subsets following transplantation.*** *Analyses of LAG3 expression on B cell subsets in B6.WT and B6.LAG3^-/-^ renal allograft recipients on d. 10 posttransplant. Quantification of frequency of LAG3 expressing cells in different B cell subsets following transplantation. The dashed line represents the mean percentage of the WT-FMO control. The data are pooled from two experiments, and each symbol represents an individual mouse. Student’s T tests were performed and p<0.05 were considered significant.*

Figure S6

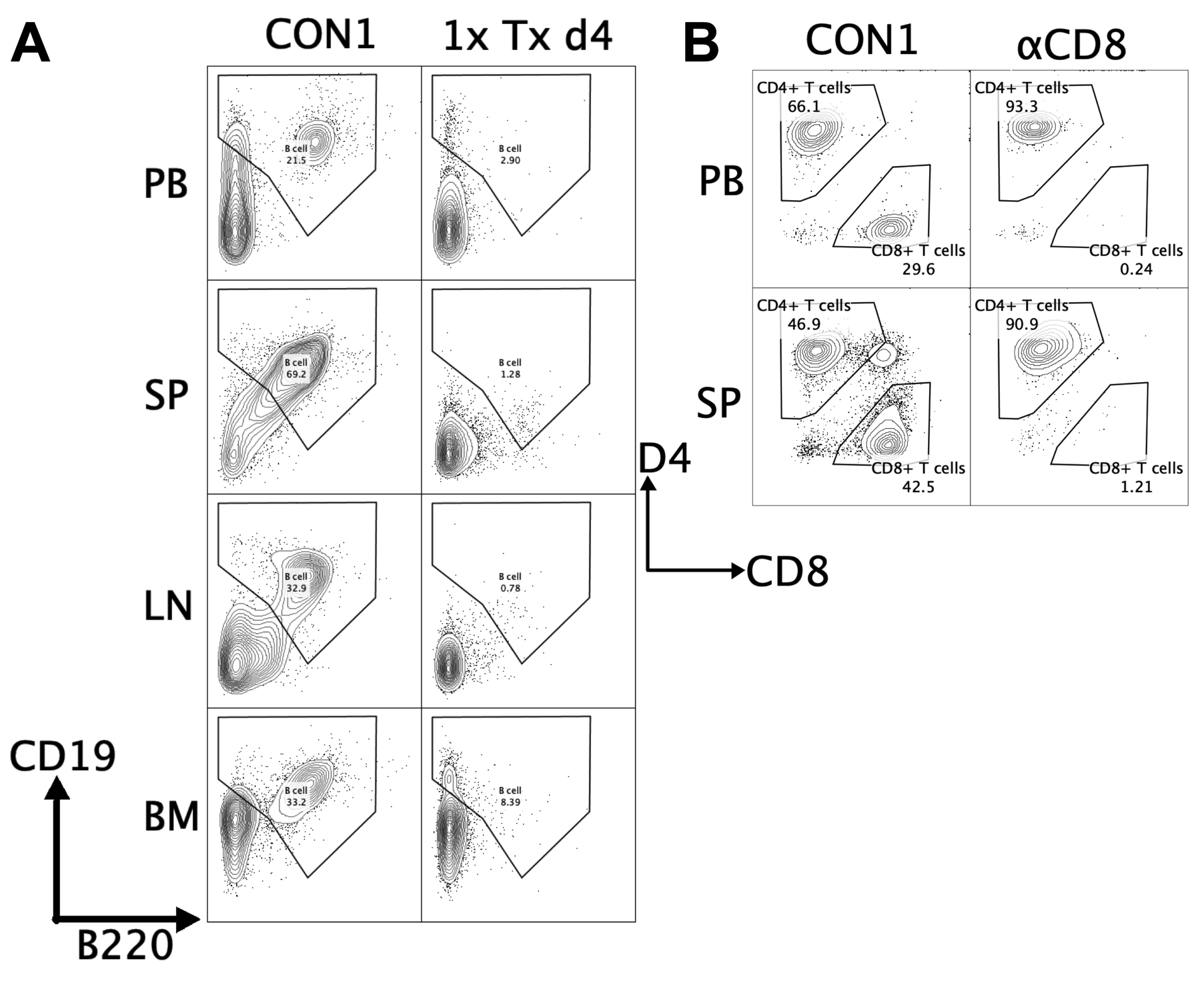

***Figure S6. Confirmation of lymphocyte depletion strategies****.* ***A****. B cells were depleted in B6.WT or B6.LAG3^-/-^ recipients after transplantation of C3H renal allografts. Peripheral blood (PB), spleen (SP), inguinal lymph nodes (LN) and bone marrow (BM) were recovered on d4 and analyzed for the presence of CD19^+^ and B220^+^ B cells.* ***B****. CD8^+^ T cells were depleted in B6.WT or B6.LAG3^-/-^ recipients after transplantation of C3H renal allografts. Peripheral blood (PB) and spleen (SP), were recovered on d4 and analyzed for the presence of CD4^+^ and CD8^+^ T cells.*

Figure S7

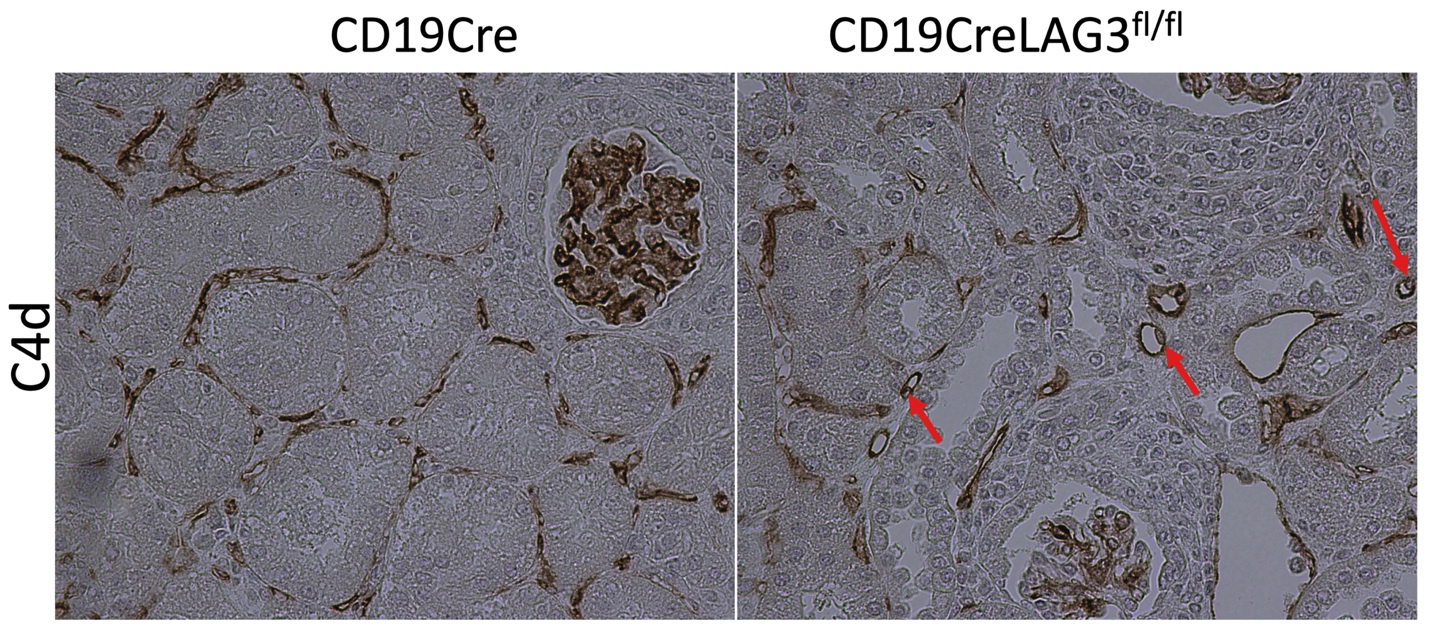

***Figure S7. Graft histology of B cell LAG3 conditional knockout mice have characteristic signs of antibody mediated rejection****. CD19Cre (left) or CD19CreLAG3^fl/fl^ (right) were transplanted with C3H kidney allografts that were recovered on d30 posttransplant and stained for C4d. Red arrows indicate swollen endothelial cells indicating antibody mediated rejection.*

Figure S8

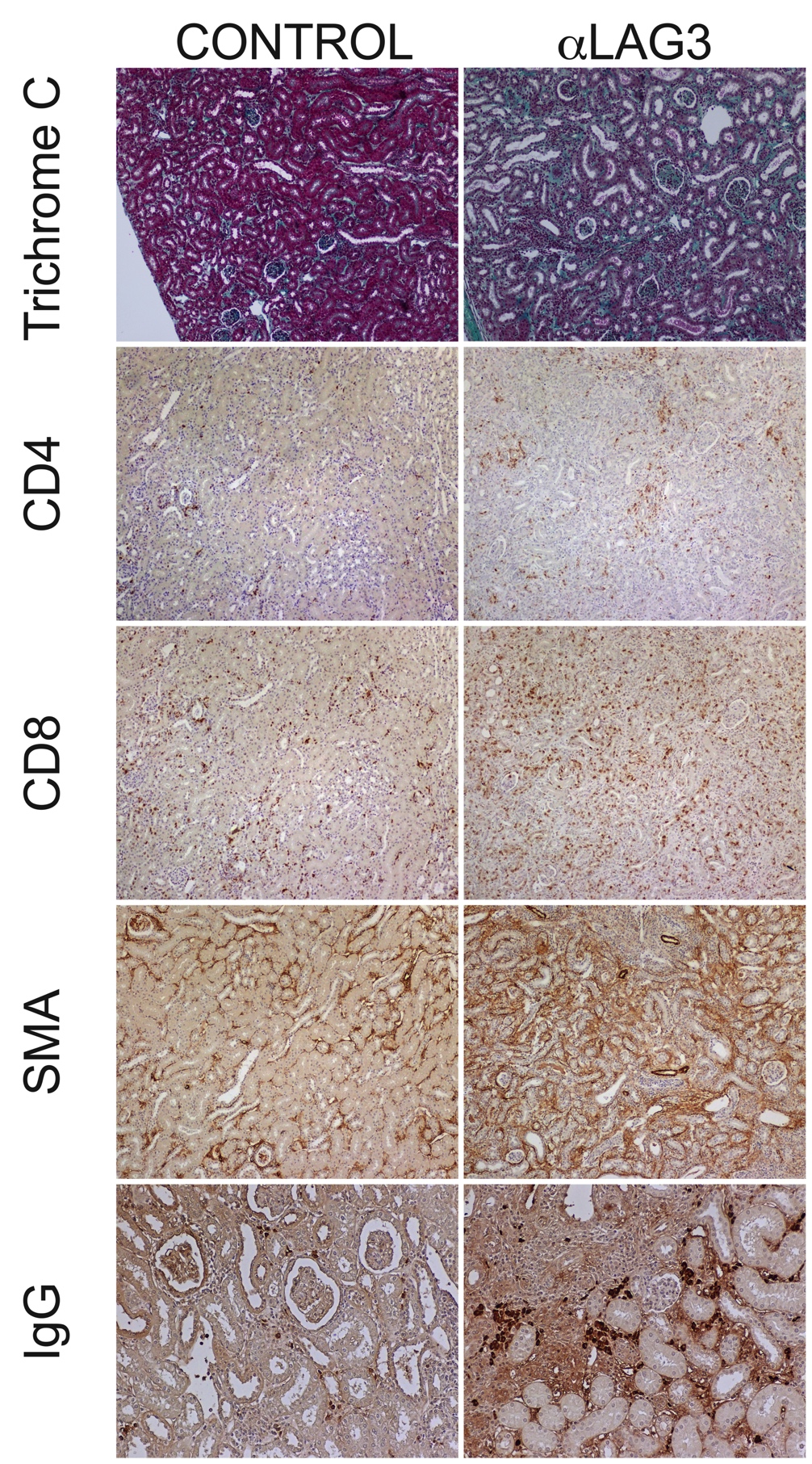

***Figure S8.*** ***LAG3 blockade following kidney transplantion leads to chronic antibody mediated graft injury.* A.** *B6.WT treated with anti-LAG3 mAb (Clone C9B7W) or control IgG posttransplantation of C3H renal allografts.Renal allografts analyzed on d. 42 posttransplant by Trichrome C and immunoperoxidase staining for CD4, CD8, smooth muscle actin (SMA) unconjugated IgG as a marker of plasma cells The photographs were taken at 200x and are representative of 4-5 animals in each group*
